# Comparative Study on Acyl Transferases in Fatty Acid and Polyketide Synthases

**DOI:** 10.1101/2020.12.29.424742

**Authors:** Franziska Stegemann, Martin Grininger

## Abstract

Fatty acid and polyketide synthases (FASs and PKSs) synthesize physiologically and pharmaceutically important products by condensation of acyl building blocks. In both multidomain enzymes, the acyl transferase (AT) is the key player responsible for the selection of these acyl units for further processing. In this study, the AT domains of different PKS systems are kinetically described in their substrate selectivity, AT–Acyl carrier protein (ACP) domain-domain interaction, and enzymatic kinetic properties. The ATs of modular PKSs, the proteins assembling the most intricate polyketides, turned out to be significantly slower than ATs from iterative FAS and PKS systems, but also more substrate specific. We explain these substantially different properties by the phylogenetically early splitting of species. For the AT of the 6-deoxyerythronolide B synthase (DEBS), the interaction with ACP is analyzed in detail by site-directed mutagenesis of interface residues. Among others, a surface exposed arginine (R850) was replaced by three different residues, leading to mutants with severely different kinetics that cannot be explained by simple effects. Our study enlarges the understanding of ATs in its molecular properties, and is similarly a call for thorough AT-centered PKS engineering strategies.

## Introduction

Fatty acid synthases (FASs) and polyketide synthases (PKSs) generate important products of primary and secondary metabolism. Palmitic and stearic acid are the main products of FASs and serve a multitude of cellular functions. ^[1]^ Polyketides are secondary metabolites of elaborate chemistry and of high bioactivity, ^[2]^ and several are harnessed for therapeutic treatment; e. g., as antibiotic (erythromycin, pikromycin, chlorothricine), immunosuppressant (rapamycin) or antineoplastic (epothilone) agent. ^[3]^ Despite the fact that they are chemically different and involved in different cellular functions, fatty acids and polyketides share a common biosynthetic origin.

Fatty acids and polyketides are assembled from small building blocks, the acyl substrates, which are loaded and processed during the biosynthesis. FASs and PKSs occur as type II systems in which every catalytic function is provided by a separate protein, and as type I systems in which the enzymatic functions are harbored on one or several polypeptide chains. ^[4]^ FASs and PKSs consist of one distinct set of protein folds whether they occur as separate enzymes or enzymatic domains. In the following, the focus will be on type I systems, which is the predominant system in eukaryotic *de novo* fatty acid synthesis and responsible for the synthesis of the plethora of elaborate polyketides. The terms FAS and PKS will refer to type I FAS and type I PKS in the following.

The FASs and a subset of PKS systems – the iterative PKSs – use one set of catalytic domains harbored on one protein complex in repeating synthetic cycles. While FASs strictly reduce the growing chain fully in every cycle, iterative PKSs generally run non-reductive cycles in which a catalytic step of one of the cycles may be omitted, a property termed cryptic coding. ^[5–7]^ In modular PKSs, several sets of catalytic domains are lined up, organized in so-called modules, each being responsible for incorporation of one extender unit. PKSs comprise a minimal set of three catalytic domains connecting the extender units: β-ketoacyl synthase (KS), acyl transferase (AT), and acyl carrier protein (ACP). Processing functions enlarge this minimal set: β-ketoacyl reductase (KR), dehydratase (DH), and β-enoyl reductase (ER). The growing acyl chain is processed with respect to the domain composition (**Figure 1A**). In modular PKSs, a huge structural and functional variety of compounds is produced dependent on module type and module assembly. ^[8], [9]^ In running vectorial synthesis and varying the enzymatic repertoire, modular PKSs harness the synthetic capacity of this protein family in diametrically different manner to FASs (**Figure 1B**).

**Figure 1:**
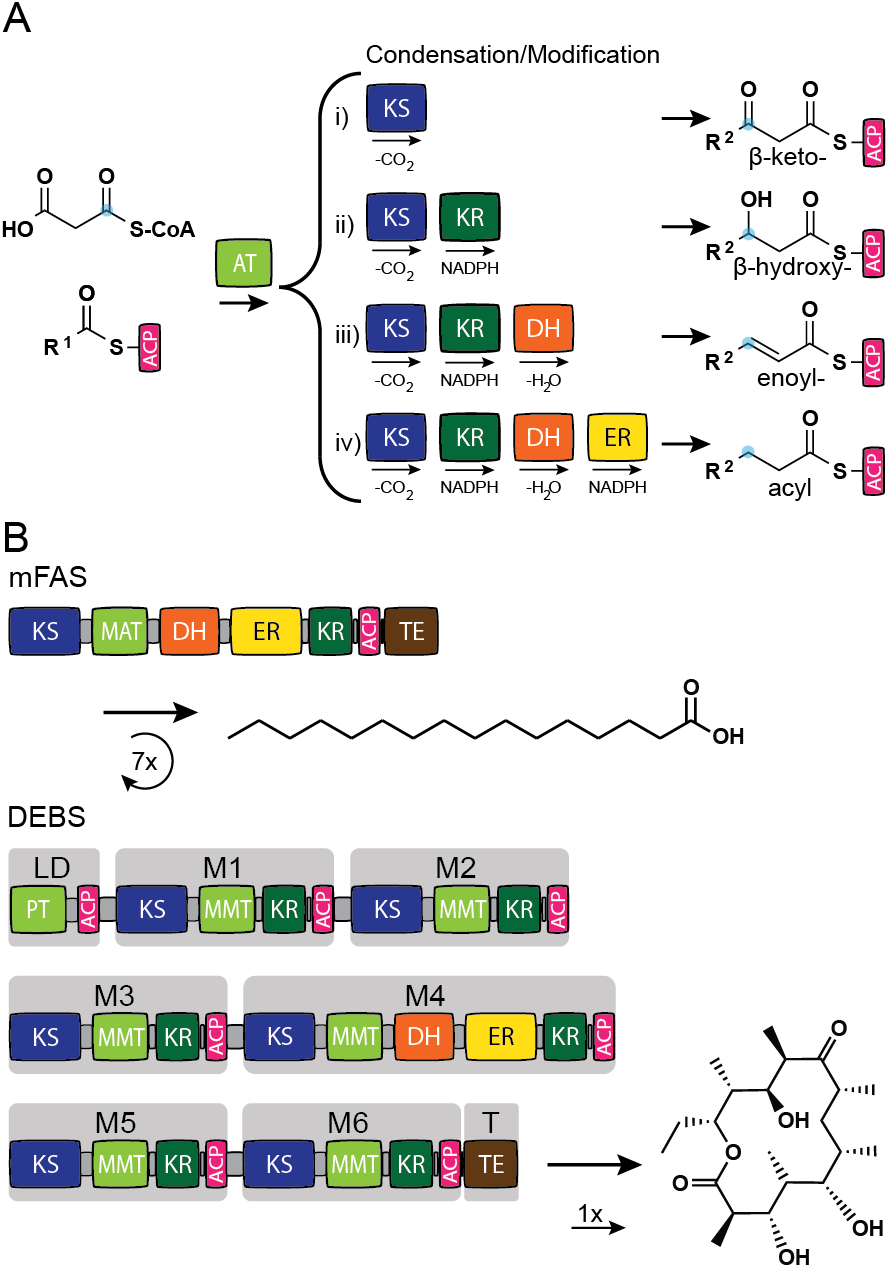
Synthesis of fatty acids and polyketides. (A) Module composition of PKSs: The minimal PKS module (i) consists of domains KS, AT, and ACP leading to a β-keto product, and can be enlarged by the modifying domains KR, DH, and ER leading to β-hydroxy (ii), enoyl (iii) or acyl (iv) products. FASs contain a full set of domains leading to fully saturated fatty acids. (B) FASs, like the murine FAS (mFAS), and iterative PKSs use their domains several times during synthesis. Modular PKSs, like 6-deoxyerythronolide B synthase (DEBS), follow an assembly-line like synthesis in which each domain is only used once during synthesis and the growing polyketide chain is passed from one module to another. Abbreviations: PT: propionyl transferase, MMT: methylmalonyl transferase, TE: thioesterase, LD: loading module, M1-M6: module 1-6, T: termination module.

The modular PKSs are one of the most elaborate proteins found in nature. The multidomain proteins evolved from the type II systems with their separated enzymes, and duplication and/or recombination events followed by more subtle adaptions further led to multimodular arrangements. ^[1], [10], [11]^ While computational analysis of the rapidly growing sequence databases allows drawing evolutionary relationships within protein families (**Figure 2 and Figure S1**), recent enzymatic analyses have also given insight into the relationship between FASs and PKSs at the functional level. Here, the domains AT and KS are most interesting, because they occur obligatorily in each protein/module. For the murine and the chicken FASs, the domains AT and KS have been characterized in considerable depth. In FASs, the AT quickly samples acyl substrates with low selectivity, while the slower KS only accepts acyl substrates for condensation without β-modification (turnover rate for AT of 100-150 s^-1^ for acetyl (Ac) and malonyl (Mal) moiety determined for chicken and murine FAS ^[12], [13]^ and 31 s^-1^ determined for condensation of Ac and Mal catalyzed by the KS of chicken FAS ^[12], [14]^). The AT and KS domains of PKSs are in general less well studied. However, the knowledge about the polyketide compound family and their biosynthetic pathways, as well as attempts to manipulate modular PKSs for the production of custom compounds imply that ATs of modular PKSs are substrate specific, ^[15–17]^ while the specificity of the KSs is less restrictive. ^[18–20]^ Unfortunately, enzyme kinetic studies on PKSs are rare, so that the above view is not underpinned by systematic and quantitative data and largely remains notional.

**Figure 2:**
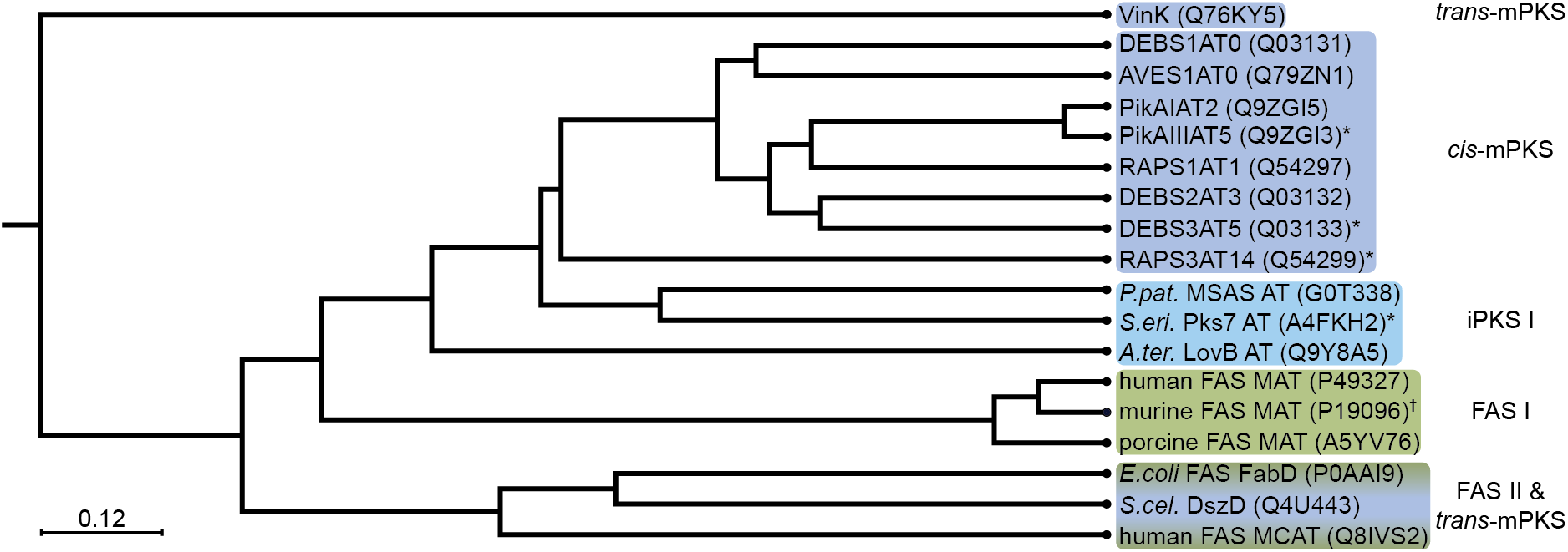
Schematic representation of the phylogenetic relationship between AT domains from FASs and PKSs. ATs from modular PKSs form a distinct clade with a subclade for the loading ATs. As expected, RAPS3 AT14 accepting Mal is separated from the MMal transferring ATs. ATs from iterative PKSs fall into a distinct clade. FAS II ATs form a distinct clade with the *trans*-acting AT DszD from disorazole PKS. All three ATs transfer Mal as substrate. The *trans*-acting AT VinK from vicenistatin PKS was expected to fall into a clade with the latter ATs ^[21]^ but is separated, which can be explained by its transfer of an unusual substrate – a dipeptidyl moiety. ^[22]^ FAS systems depicted in green, PKS systems depicted in blue. Type I systems shown in light, type II systems shown in dark colors. ATs analyzed in this study are indicated by *. AT analyzed in a previous study and discussed in this study is marked with ^†^. ^[13]^ Abbreviations: iPKS: iterative PKS, mPKS: modular PKS.

In this study, we kinetically describe the AT domain in order to understand similarities and differences in its function in the context of the different megasynthase systems. Three different modular PKSs – one incorporating Mal, two incorporating methylmalonyl (MMal) – are compared to an iterative PKS and to type I and type II FAS systems. We collect a large body of enzyme kinetic data, and are eventually able to give insight into the function of ATs in FASs and PKSs: (i) Our data proposes that ATs of PKSs are in general slower than of FASs, which may reflect the different scopes of biosynthetic pathways. Whereas fatty acids are essential compounds of all living entities (except archaea) and needed in high amounts, polyketides are secondary metabolites that can take effect at low concentrations. (ii) We observe in our dataset that the slower ATs of the modular PKSs do not load non-native substrates even in absence of the native substrates, which is different to the faster iterative ATs, indicating high substrate specificity for the modular ATs only. We hypothesize that during the evolutionary development of modular PKSs from genes and gene sequences by duplication, recombination and/or gene conversion, ATs of modular PKSs have developed substrate specificity to compensate for the relaxed specificity of KS-mediated condensation step. (iii) We further see that both steps in the two-step (ping-pong) mechanism of transacylation contribute to substrate specificity, and that this selectivity can emerge in either the first or the second step.

Furthermore, we analyze the interaction between AT and ACP in a mutational study on module 5 (M5) of the modular PKS 6-deoxyerythronolide B synthase (DEBS). We model the AT:ACP interface by homology to interfaces recently solved in structure, introduce point mutations in the AT domain, and record transacylation rates. Our data shows that surface mutations modulate transacylation rates in intricate manner, speaking against high success rates of oversimplified engineering strategies. Particularly, the surface amino acid R850 of DEBS M5 is a hot spot in this regard.

## Results and Discussion

### Kinetic Analysis of AT-Mediated Reactions

The AT domain catalyzes the transfer of the acyl moiety X of the substrate X-coenzyme A (CoA) onto the *holo*-ACP domain (**Figure 3)**. The transacylation reaction follows a double displacement mechanism, called Ping-Pong Bi Bi mechanism. ^[23]^ The ping step denotes the first transfer of the acyl moiety X onto the enzyme AT, resulting in release of product CoA. The pong step denotes the second transfer of the acyl moiety X onto the ACP after binding of the second substrate ACP to AT-X, leading to the final product ACP-X. Hydrolysis occurs as side reaction, releasing the free acid “X-OH”.

**Figure 3:**
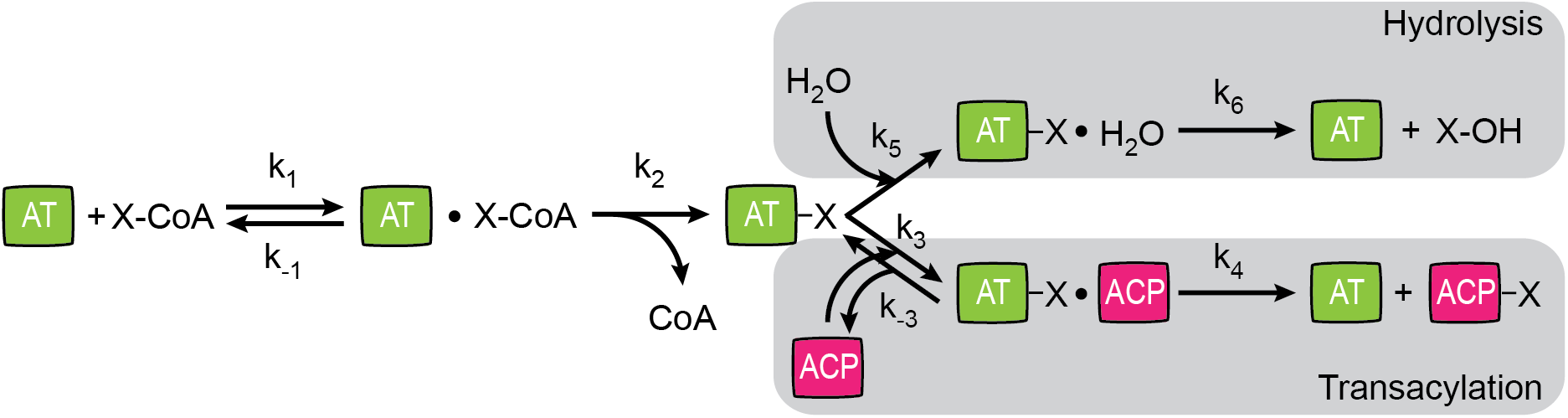
Schematic representation of AT-mediated reactions. After formation of the enzyme (AT)-substrate (X-CoA) complex, free CoA is released and AT-X is formed. The substrate X is either hydrolyzed during AT-mediated hydrolysis (upper branch) or transacylated via formation of the enzyme (AT-X)-substrate (ACP) complex (lower branch). Kinetic constants *k_1_* to *k_6_* describe the AT-mediated reactions.

To quantify transferase activity, an enzyme-coupled, fluorescence-based assay was established, based on previous work. ^[13], [16], [24]^ It couples formation of free CoA with the reduction of nicotinamide adenine dinucleotide (NAD^+^), which can then be followed fluorimetrically, allowing the quantification of transferase activity. The acyl transfer is measured in presence and absence of ACP, giving access to transacylation and hydrolysis rates, respectively. Kinetic hydrolysis parameters are calculated by titration of X-CoA. To determine absolute kinetic transacylation parameters, X-CoA is titrated to a series of fixed ACP concentrations. Kinetic analysis of transacylation and hydrolysis gives the following parameters arising from the kinetic constants *k_1_* to *k_6_*. Notably,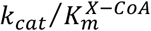 is the same for transacylation and hydrolysis, as seen in these mathematical derivations for transacylation and hydrolysis (**Figure 4**). For details, see **Supporting Information Note 1**.

**Figure 4:**
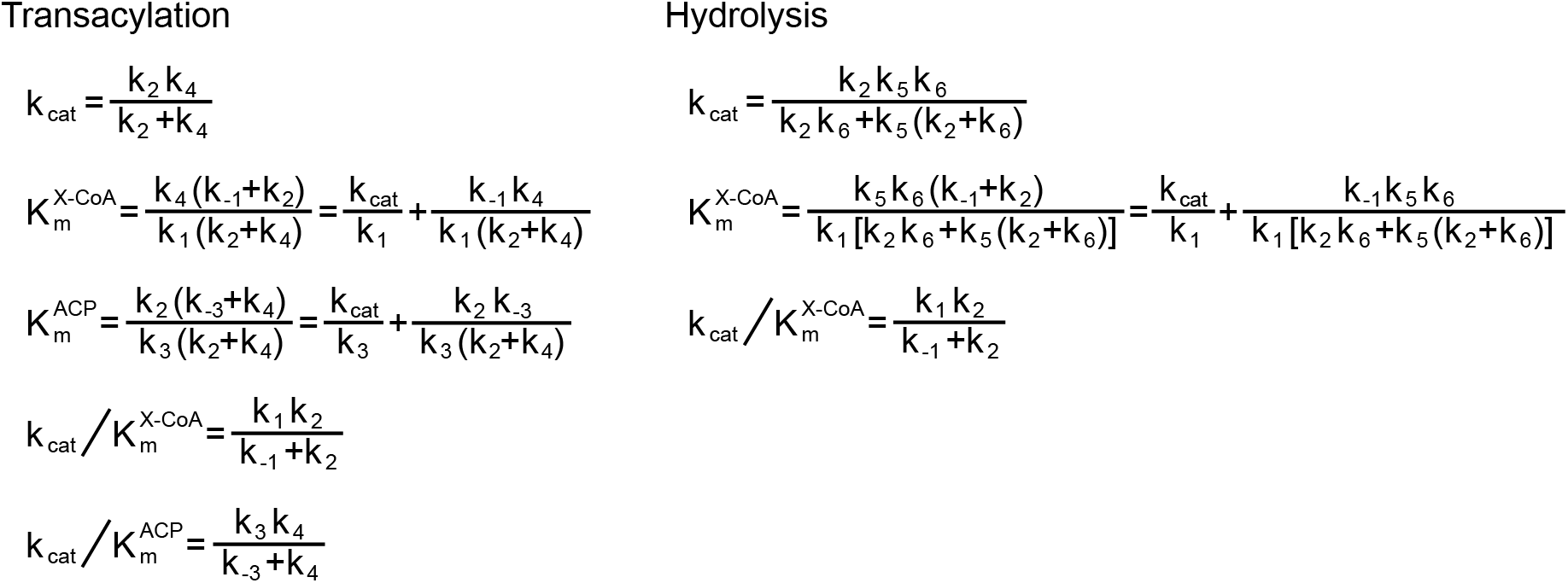
Kinetic parameters defined by kinetic constants *k_1_* to *k_6_* describing the AT-mediated reactions – transacylation and hydrolysis. The turnover number *k_cat_* and the Michaelis-Menten constants 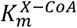 and 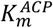 for both substrates as well as the corresponding catalytic efficiencies 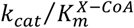 and 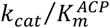.

### Expression and Purification of AT and ACP Domains

In order to understand similarities and differences between types of FASs and PKSs, the following AT domains of some representatives were analyzed in this study: Representing the AT domain of modular PKSs, the AT of DEBS M5 from *Saccharopolyspora erythraea* (termed DEBS3M5), of pikromycin synthase (Pik A) M5 from *Streptomyces venezuelae* (PikAIIIM5), and of rapamycin synthase (RAPS) module 14 from *Streptomyces hygroscopicus* (RAPS3M14) were used. The first two AT domains transfer MMal, while the latter transfers Mal. ^[2], [25]^ For the iterative PKSs, we worked with the AT domain of Pks7 found in *Saccharopolyspora erythraea*. Pks7 AT is phylogenetically similar to 6-methylsalicylic acid synthase (MSAS) AT from *Penicillium patulum*, and is likely involved in methylsalicylic acid (MSA) synthesis (**Figure 2**). ^[26]^ Data will be compared to the previously analyzed murine FAS (mFAS), which was found to be promiscuous in transferring a large variety of substrates. ^[13]^ The AT domain of Pks7 is expected to load Ac and Mal onto the ACP domain for MSA production. ^[3], [7], [27]^ The specificity for Mal, and not MMal, is supported by the amino acid fingerprint of Pks7 AT. ^[3]^

Recent functional and structural characterization has revealed that FAS and PKS systems form stable KS-AT dimers. ^[13], [28]^ Therefore, all AT domains used in this study were expressed as KS-AT didomains. A KS knockout, leading to constructs denoted as KS^0^-AT, allowed for selectively inspecting the AT domain in transacylation properties. Recombinant expression in *E. coli* gave sufficient yields of soluble proteins (**Table S1)**, which were pure after tandem purification as judged from sodium dodecyl sulfate polyacrylamide gel electrophoresis (SDS-PAGE) (**Figure S2**). Size exclusion chromatography (SEC) (**Figure S3**) revealed stable dimers for Pks7 and PikAIIIM5 KS^0^-AT, and tetrameric and dimeric oligomers for DEBS3M5 KS^0^-AT. Surprisingly, RAPS3M14 KS^0^-AT eluted from SEC mainly as monomer, besides a subfraction of tetrameric species. We note that the prevalence for a monomeric state should not impact AT activity, since the dimeric interface is formed by the KS domain, while the AT domain is monomeric in its active form. A thermal shift assay (TSA) was performed in two buffer systems to validate protein quality, giving melting points within a range of 45-51 °C and 36-42 °C in storage and assay buffer, respectively (**Table S2 and Figure S4**). We note that all AT domains were prepared in biological triplicates, produced from single clones of one or more independent plasmid transformations.

Complete kinetic characterization of AT domains requires high yield and high quality of the corresponding ACPs. In order to generate *holo*-ACP, co-expression of ACPs with the 4’-phosphopantetheinyl transferase from *Bacillus subtilis* (Sfp) was performed, which, however, resulted in insufficient yields of most ACPs. Using codon-optimized sequences, the *holo*-ACP yields increases significantly. ACPs were purified by affinity chromatography and SEC of which the latter allowed selecting for soluble ACPs. For typical expression yields and SDS-PAGE analyses, see **Table S3 and Figure S2**. Quantitative *holo*-ACP formation was confirmed by mass spectrometry (**Table S4 and Figure S5**). Protein integrity was confirmed by TSA giving melting temperatures within a range of 57-64 °C and 50-65 °C in storage and assay buffer, respectively (**Table S2 and Figure S6**).

### Screening of Substrates in AT-mediated Reactions

At the outset of the functional characterization of AT domains, we performed an initial substrate screen. In doing so, three substrates were used: Mal and MMal, as typical extender units, and Ac, as typical priming unit, and likewise a negative control for the ATs of modular PKSs. For the initial substrate screen, AT domains were probed at fixed ACP and X-CoA concentrations.

With our initial screen, we essentially confirmed the substrate specificity of the modular PKSs (DEBS3M5, PikAIIIM5 and RAPS3M14) as expected from the chemical structure of the product (**Figure 5A, and Table S6**): The AT domains of DEBS3M5 and PikAIIIM5 transferred solely MMal, and RAPS3M14 solely Mal. The AT domains of the iterative Pks7 tolerated MMal besides its natural extender substrate Mal, as well as the priming substrate Ac at slower rate. Since MMal-CoA is present in the native host *S. erythraea*, the loose specificity implies that both extender substrates can be loaded for condensation. We analyzed Pks7 in transacylation of both substrates for deeper insight in the following (see below). For the MAT domain of mFAS (mMAT), broad substrate tolerance was already shown previously, including the fast transfer of the priming substrate Ac and extender substrate Mal with similar rates as well as the transfer of MMal at lower rate. ^[13]^

**Figure 5:**
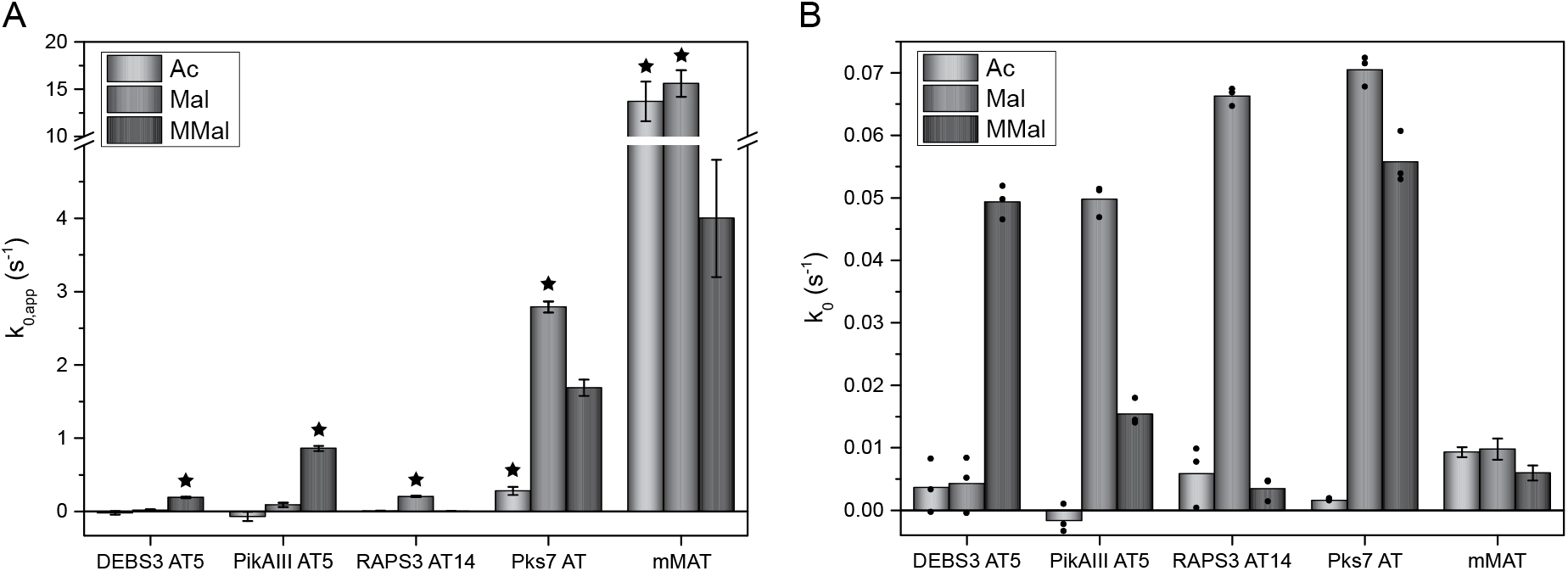
Initial screening of FAS and PKS systems. (A) Transacylation screening: loading of Ac-(light grey), Mal-(grey) and MMal-CoA (dark grey) is measured in presence of ACP. Substrate concentrations (X-CoA and ACP) are fixed at 50 μM. Bars indicate the average substrate turnover. A star indicates the native substrates for transacylation. Error bars are calculated for biological triplicates. mMAT data from a previous study, referring to *k_cat,app_* at saturated X-CoA concentration and 60 μM ACP. ^[13]^ (B) Hydrolysis screening. Loading of Ac-(light grey), Mal-(grey) and MMal-CoA (dark grey) is measured in absence of ACP. Substrate concentration (X-CoA) is fixed at 50 μM. Bars indicate the average substrate turnover. Black dots correspond to the average of technical triplicates of one biological replicate. mMAT data from a previous study, referring to *k_cat_* at saturated X-CoA concentration. ^[13]^

Probing AT-mediated hydrolysis, we observed that most domains transfer substrates with the same preference in hydrolysis and in transacylation (**Figure 5B**). The AT domain of PikAIIIM5 is an exception, in showing different preferences in hydrolysis and transacylation. Overlapping substrate specificity of transacylation and hydrolysis has also been reported in a previous study for the AT domain of DEBS module 3 (DEBS2M3). ^[16]^ This data rules out that hydrolysis is a proofreading function for clearing the active site from wrongly loaded acyl moieties, as previously suggested. ^[29], [30]^

We note, that for determination of transacylation rates, self-acylation of ACP and not AT-mediated hydrolysis was subtracted as background. Accordingly, in this initial screening, the PKSs’ transacylation rates may contain hydrolysis rates to some extent.

### Kinetic Characterization of Hydrolysis

Based on the initial screening, full kinetic analysis of AT-mediated hydrolysis and transacylation was performed with the substrates that are preferentially transferred. In order to analyze AT-mediated hydrolysis, the substrates X-CoA were titrated in absence of ACP.

Data collection for recording hydrolysis suffered from extreme differences in 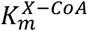, and we were just able to determine kinetic parameters precisely for 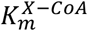 values lying within suited conditions for recording Michaelis-Menten kinetics. (i) For high Michaelis-Menten constants, high substrate concentrations have to be used. This can lead to experimental problems caused for example by substrate inhibition effects, and only rough values for 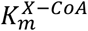 and *k_cat_* can be given, leading also to catalytic efficiencies 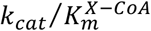 that are error-prone. (ii) For low Michaelis-Menten constants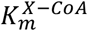, low substrate concentrations have to be used, causing poor signal-to-noise ratios in the assay. Increasing enzyme concentrations lead to disregard of Michaelis-Menten concentration requirement. Thus, for AT domains with low 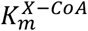, just *k_cat_* is determined precisely, whereas 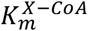 and therefore also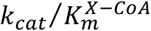 are approximate values.

Given these constraints, only DEBS3M5 AT-mediated hydrolysis was eventually characterized in 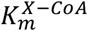 and *k_cat_* at high confidence, while for the other AT domains of PKSs partly just rough values were determined (**Table 1**). The kinetic parameters for DEBS3M5 AT-mediated hydrolysis are very similar to the data for DEBS2M3 AT-mediated hydrolysis previously published by Dunn *et al.* ^[16]^ Overall, AT-mediated hydrolysis rates for the native substrates are similar for all analyzed PKS systems. Hydrolysis mediated by mMAT is very slow compared to the PKS systems. For hydrolysis titration curves, see **Figure S7**.

**Table 1.** Kinetic parameters determined for AT-mediated hydrolysis. Abbreviations: n.d.: not determinable.

| Protein<br>X-CoA | $k_{cat}$ ($s^{-1}$) | $K_m$, X-CoA ($\mu M$) | $k_{cat}/K_m$, X-CoA<br>($M^{-1}s^{-1}$) |
| --- | --- | --- | --- |
| DEBS3AT5<br>MMal | $5.31 \times 10^{-2} \pm 9.62 \times 10^{-4}$ | $7.88 \pm 3.98 \times 10^{-1}$ | $6.7 \times 10^3$ |
| PikAIIIAT5<br>MMal [a] | $1.94 \times 10^{-2} \pm 5.03 \times 10^{-4}$ | $<8.91 \times 10^{-1}$ | $>2.2 \times 10^4$ |
| PikAIIIAT5<br>Mal [b] | $>1.95 \times 10^{-1}$ | $>100$ | n.d. |
| RAPS3AT14<br>Mal [a] | $6.51 \times 10^{-2} \pm 1.89 \times 10^{-3}$ | $<5.45 \times 10^{-1}$ | $>1.2 \times 10^5$ |
| Pks7 AT<br>Mal [a] | $9.93 \times 10^{-2} \pm 3.17 \times 10^{-3}$ | $<4.74 \times 10^{-1}$ | $>2.1 \times 10^5$ |
| Pks7 AT<br>MMal<sup>[a]</sup> | $7.34 \times 10^{-2} \pm 1.51 \times 10^{-3}$ | $<3.98 \times 10^{-1}$ | $>1.8 \times 10^5$ |
| mMAT<br>Mal<sup>[c]</sup> | $(9.8 \pm 1.7) \times 10^{-3}$ | n.d. | n.d. |
| mMAT<br>MMal<sup>[c]</sup> | $(6.0 \pm 1.2) \times 10^{-3}$ | n.d. | n.d. |
| mMAT<br>Ac<sup>[c]</sup> | $9.3 \times 10^{-3} \pm 1.2 \times 10^{-4}$ | n.d. | n.d. |

[a] This system has a very low $K_m^{X-CoA}$ . Therefore, Only $k_{cat}$ is determined precisely. [b] This system has a very high $K_m^{X-CoA}$ . Thus, kinetic parameters determined are only minimal values. [c] mMAT data from previous study. <sup>[13]</sup>

### Kinetic Characterization of Transacylation

In order to characterize the transacylation reaction, X-CoA substrates were titrated to a series of fixed ACP concentrations. For all ATs, absolute kinetic parameters were received, although the kinetic parameters for systems with high 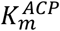 could be determined less accurately, simply because the vast amounts of ACP needed to cover an appropriate substrate range could not always be provided. This was particularly problematic for Pks7. Here, the ACP concentration was varied only within a range of 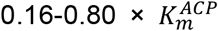 for the substrate MMal-CoA.

In the following, we compare the tested systems in turnover rates, Michaelis-Menten constants, and catalytic efficiencies. Overall, data reveals that PKS systems feature slower turnover rates than the FAS type I and type II systems (**Table 2 and Figure 6)**, ^[13], [31], [32]^ with the AT domains of iterative Pks7 transferring the substrates with significantly higher rates than AT domains of modular PKSs. Catalytic efficiencies 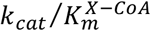 and 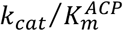 show that the mMAT domain transacylates with highest efficiency, followed by the iterative Pks7 AT (about one order of magnitude less efficient than mMAT) and modular PKSs (another order of magnitude less efficient than Pks7 AT), indicating that the iterative systems feature energetically low transition states for the transacylation reaction. Particularly, the catalytic efficiencies of the AT-acylating ping step 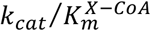 is outstandingly high for mFAS compared to modular PKSs; e. g., Mal is transferred by mMAT with catalytic efficiency of 8.3 × 10^6^ M^-1^s^-1^ compared to RAPS3 AT14 with 4.8 × 10^4^ M^-1^s^-1^. Comparing the ratio of the catalytic efficiencies, we note, that the iterative systems show 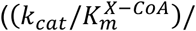 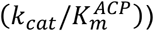of around 30 for native extender substrates. For the slower modular PKS systems, this ratio drops significantly, and in case of DEBS3 AT5, it is inverted. The ratio of catalytic efficiencies may be key to high turnover rates in iterative systems: The high efficiency of the ping step leads to more acylated AT. Since the high AT-X concentrations increase the chance for productive AT-X–ACP interactions, i. e. the successful loading of ACP, higher turnover rates will be achieved. Vice versa, a lower efficiency in the ping step likewise promote frequent non-productive interactions, which then slows down the transfer of the acyl moiety onto ACP.

**Table 2.** Kinetic parameters determined for transacylation.

| Protein<br>X-CoA | $k_{cat}$ ($s^{-1}$) | $K_m$, X-CoA<br>($\mu M$) | $K_m$, ACP ($\mu M$) | $k_{cat}/K_m$, X-CoA<br>($M^{-1}s^{-1}$) | $k_{cat}/K_m$, ACP /<br>($M^{-1}s^{-1}$) | $k_{cat}$ norm<br>(%) | $k_{cat}/K_m$, X-CoA<br>norm (%) | $k_{cat}/K_m$, ACP<br>norm (%) | $k_{cat} /$<br>$K_m$, X-CoA /<br>($k_{cat} /$<br>$K_m$, ACP) |
| --- | --- | --- | --- | --- | --- | --- | --- | --- | --- |
| DEBS3AT5<br>MMal | $1.25 \pm 7.11 \times 10^{-2}$ | $209 \pm 17.5$ | $72.0 \pm 8.21$ | $6.0 \times 10^3$ | $1.7 \times 10^4$ | 1.1 | $4.3 \times 10^{-2}$ | 3.5 | $3.4 \times 10^{-1}$ |
| PikAIIIT5<br>MMal | $7.47 \pm 5.16 \times 10^{-1}$ | $96.2 \pm 8.69$ | $330 \pm 44.1$ | $7.8 \times 10^4$ | $2.2 \times 10^4$ | 6.3 | $5.6 \times 10^{-1}$ | 4.5 | 3.4 |
| RAPS3AT14<br>Mal | $7.10 \times 10^{-1} \pm 3.84 \times 10^{-2}$ | $14.8 \pm 1.33$ | $66.0 \pm 10.8$ | $4.8 \times 10^4$ | $1.1 \times 10^4$ | $6.0 \times 10^{-1}$ | $3.4 \times 10^{-1}$ | 2.2 | 4.6 |
| Pks7 AT<br>Mal | $23.4 \pm 1.50$ | $9.48 \pm 8.84 \times 10^{-1}$ | $286 \pm 30.3$ | $2.5 \times 10^6$ | $8.2 \times 10^4$ | 20 | 18 | 17 | 30 |
| Pks7 AT<br>MMal | $20.5 \pm 2.03$ | $21.1 \pm 2.46$ | $505 \pm 73.8$ | $9.7 \times 10^5$ | $4.1 \times 10^4$ | 17 | 6.9 | 8.4 | 24 |
| mMAT<br>Mal [a] | 119 | 8.8 | 245 | $1.4 \times 10^7$ | $4.9 \times 10^5$ | 100 | 100 | 100 | 28 |
| mMAT<br>Ac [a] | 99.2 | 12 | 265 | $8.3 \times 10^6$ | $3.7 \times 10^5$ | 83 | 59 | 76 | 22 |

[a] mMAT data from previous study. [13]

**Figure 6:**
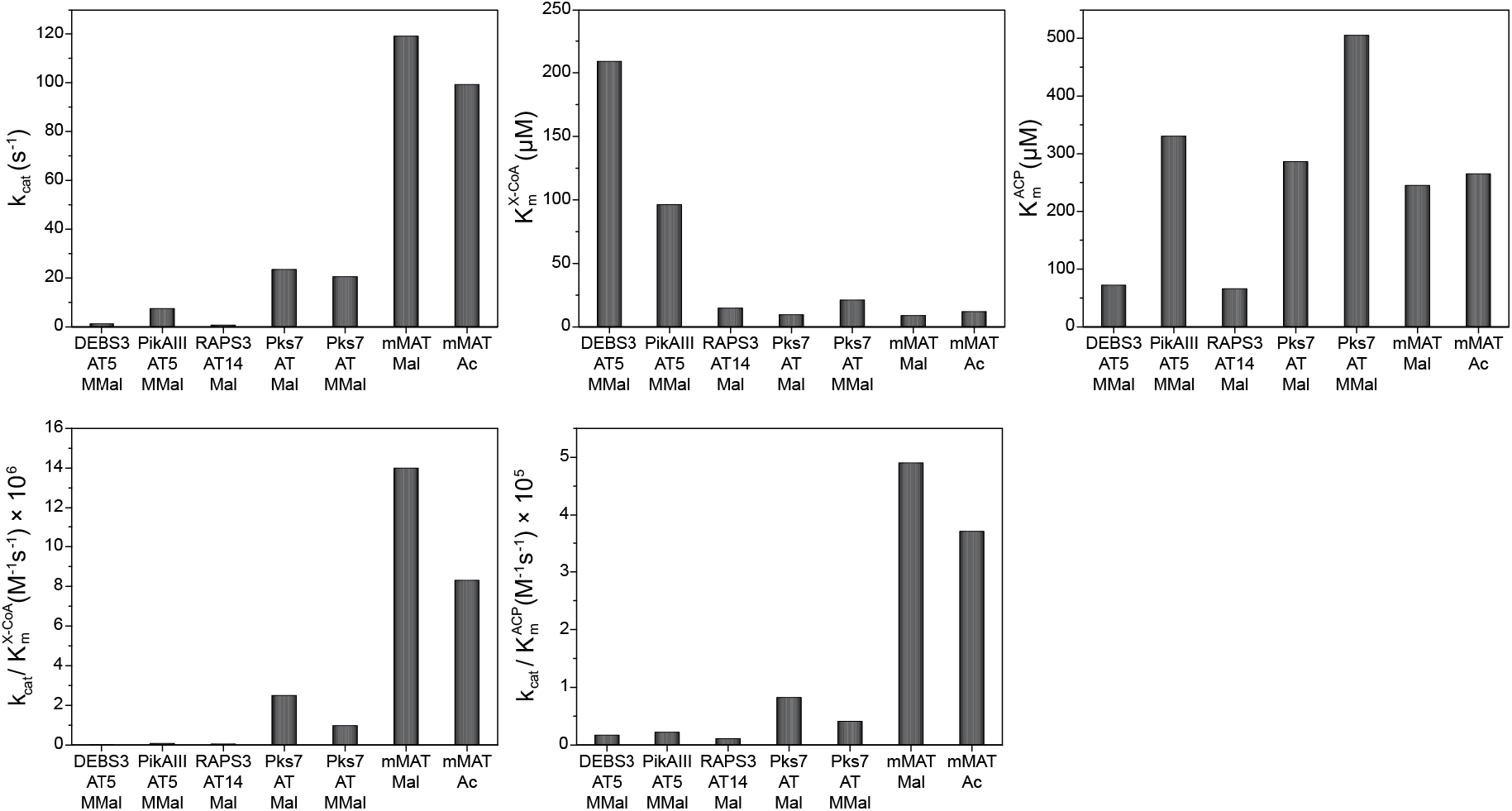
Turnover rate, Michaelis-Menten constants and catalytic efficiencies determined for transacylation for the FAS and PKS systems. mMAT data from previous study. ^[13]^

The 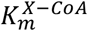 values for Ac-CoA and Mal-CoA are in the range of bacterial metabolite levels, ^[33]^ and are similar for most AT substrate pairs. Interestingly, the 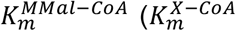 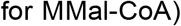values for DEBS3 AT5 and PikAIII AT5 are 10 to 20-fold higher than 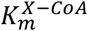 for other systems tested. This either indicates an adaptation to higher cellular MMal-CoA levels or a control mechanism, regulating the MMal turnover depending on its availability. Due to the high 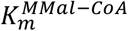, modular PKSs operate at low turnover rates for low MMal-CoA concentrations, so that the metabolic flux into the polyketide pathway is limited for shortage of MMal-CoA. Higher MMal-CoA concentrations could likewise increase turnover rates of modular PKSs and the secondary metabolite output. We note that a similar regulation of polyketide synthesis would not be plausible with Mal-CoA, which is dedicated mainly to the central metabolism of any bacterial cell, and regulated in concentration to its need as precursor for fatty acid biosynthesis. ^[34]^

Our data further shows that Michaelis-Menten constants 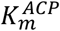 vary moderately for the different ATs. Considering molar concentrations within the FAS/PKS compartment of about 1 mM (rough calculation based on a cylinder volume of dimensions taken from MSA-like PKS; i. e., radius 10 nm, height 15 nm; 2 molecules) as well as freely diffusing domains within the multidomain compartment, all AT domains of type I systems operate at high (Pks7) to saturated (other systems tested) ACP conditions so that the pong step runs at maximum rate (*v_max_*). For the *E. coli* type II FAS, 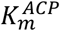 values from 40 μM to 351 μM have been reported, which is within the range of values recorded by us for the type I systems. ^[32], [35], [36]^ For *E. coli* type II FAS, 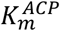 values are higher than cellular ACP concentrations of 4.6 to 14 μM (calculated from *E. coli* ACP (AcpP) copy numbers reported by Ishihama *et al.* ^[37]^ and an average volume of an *E. coli* cell of 2.8 μm^3^), ^[38], [39]^ which implies the *E. coli* AT (FabD) operates below *v_max_* and the rate is sensitive to variations in AcpP concentration.

Regarding substrate selectivity processes in iterative PKSs, Pks7 AT shows interesting features: Both Michaelis-Menten constants 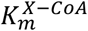 and 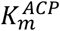 differ for the two tested extender substrates Mal-CoA and MMal-CoA. Regarding the catalytic efficiency of both steps, ping step and pong step are more efficient for Mal, which allows double selection for the native over the non-native substrate in presence of both. Assuming a lower cellular concentration of MMal-CoA than of Mal-CoA, specific condensation with Mal is likely ensured. Differences in 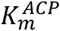 further imply that the interface between AT and ACP is responding to the substrate to be transferred, similarly as recently suggested for *E. coli* type II FAS. ^[31]^ For transacylation titration curves, see **Figure S8**.

### Further Implications from Kinetic Analysis

Comparing hydrolysis and transacylation, we observe that hydrolysis mediated by ATs of PKS systems is just one to two orders of magnitude slower than transacylation. This rate difference is much less than for FAS systems for which we had recorded four orders of magnitude difference. ^[13]^ We observe AT-mediated hydrolytic activity for all substrates – if native or non-native – transferred onto the ACP during transacylation, and in most cases hydrolysis is fastest for the native substrates (**Figure 5A and B**). The purpose of the relatively high hydrolysis rates for PKSs remains elusive, but our data rules out any quality control by hydrolysis, as e. g. reported for some aminoacyl-tRNA synthetases in which hydrolytic active sites evolved in addition to their catalytic active sites. ^[40]^ We note that comparing the ability of ATs for transacylation and hydrolysis can give insight in the substrate specificity of enzymes. (i) PikAIII AT5 is the only enzyme from our set of proteins for which we recorded a higher hydrolysis rate for the non-native substrate, which is, however, still significantly lower than the transacylation rate for the native substrate, arguing against a proofreading function. Data implies that both substrates pass the ping step and are loaded onto the AT domain, but then follow hydrolysis/transacylation to a different extent: For the native substrate, transacylation is the predominant reaction. For the non-native substrate, hydrolysis becomes more important and the ratio between hydrolysis and transacylation rate is significantly increased. In this regard, PikAIII seems capable of differentiating between the two substrates in the pong step. (ii) In case of the other modular PKSs, the absence of hydrolytic activity for non-native substrates indicates a more stringent selection for substrates in the ping step.

We note that, based on the catalytic efficiency of the ping step, we can calculate the Michaelis-Menten constants for the AT-mediated hydrolysis (**Table S6**). This is possible, because the efficiencies for the ping step in transacylation and for hydrolysis are identical (**Figure 4**). We observe decreasing 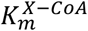 values for hydrolysis from modular PKSs to the iterative PKS to the FAS system. This trend, as well as the trend in catalytic efficiencies (increased for iterative proteins), is overall similar to the ping step of transacylation.

### Implications of Kinetic Data on FAS/PKS Function and Evolution

Bar-Even and Noor *et al.* ^[41]^ have mined enzyme databases for a systematic study on kinetic parameters of enzymes and substrates. As one of their key findings, they revealed that enzyme substrate pairs associated with primary metabolism tend to have higher turnover numbers than pairs of secondary metabolism, and also average *k_cat_/k_m_* values are higher in primary than in secondary metabolism. We observe a very similar trend in our study on the transacylation function in FASs and PKSs. Along the lines of Bar-Even and Noor *et al.*, we propose that kinetic differences result from a divergent evolutionary development of the transacylating domains (MAT and AT) in the two systems. Starting from a substrate promiscuous ancestor enzyme, the AT domain of PKSs has evolved towards higher specificity by positive selection leading to neofunctionalization via duplication and selection events. Kinetic parameters have changed towards essentially lower turnover numbers in the context of the purpose of PKSs, that is providing sufficient efficiency to produce molecules at a low basal level or when necessary without draining vital metabolic pools (particularly in case of using Mal-CoA as extender substrate). In contrast, the MAT domain of FAS was under selection pressure for providing fast rates for the high metabolic fluxes that are required by the central metabolic fatty acid biosynthesis. The phylogenetic analysis shows that the MAT is less evolved than AT domains of PKSs, and close to a FAS/PKS common precursor (**Figure 2**). We could recently demonstrate the high substrate promiscuity of mMAT, ^[13]^ which seems to be reminiscent of a primordial enzyme with this property. MAT’s substrate promiscuity may have been preserved, since in FASs the KS-mediated condensation step is highly substrate-specific, ensuring the fidelity of the fatty acid biosynthesis. ^[42]^

The molecular basis for the different substrate specificities of MAT vs. AT and their different catalytic efficiencies is not entirely clear. For the mMAT domain, we recently posited that the high conformational variability of the linkers between the α/β-hydrolase fold and the ferredoxin-like fold, as well as of the active site arginine that coordinates the free carboxylic acid of extender substrates, allows the accommodation of chemically and structurally diverse CoA-esters. ^[42]^ It would be interesting to see whether there is a more rigid geometry of the AT domains of PKSs responsible for their constrained substrate specificity. However, coverage with structural and computational data is too poor for any deeper analysis. We note that a putatively more constrained conformational variability of PKS ATs may well account for a narrow substrate specificity, but not necessarily for slow rates. Conformational variability of substrates can reduce turnover rates, when sampling states that are non-productive to catalysis. Likewise, reduced conformational variability may offer high rates for the native substrate, when sampling conformations of the catalytic cycle. ^[43], [44]^ In that light, a molecular basis for the difference in transacylation rates cannot be given.

### AT:ACP Interface Mutation Study

The incorporation of non-native extender substrates to polyketides has previously been achieved by exchanging AT domains; i. e., replacing the native domain with a domain of interest. ^[45–47]^ While such strategies can be successful in delivering the desired compound, the chimeric PKSs inherently suffer from non-cognate (permanent) interfaces in the KS-AT didomain as well as non-cognate (transient) interfaces during substrate shuttling (AT:ACP), which generally leads to compromised activity. ^[48]^ A better knowledge on substrate shuttling may offer the chance of adapting a non-cognate AT:ACP interface to kinetic properties of the overall system. Some of the molecular details to the ACP-mediated shuttling of substrates and intermediates to the catalytic domains in FASs and PKSs are being currently revealed. For example, interactions are guided by weak electrostatic interactions, ^[49]^ and, as shown for the AT–ACP interaction in the *E. coli* type II system (FabD:AcpP interface), do not necessarily involve prevalent interfaces. ^[36]^ However, in spite of this progress, substrate shuttling remains poorly understood, which hinders efficient PKS engineering. ^[50]^

In order to understand this critical domain-domain interaction and guide future PKS engineering strategies, the AT:ACP interface of DEBS3M5 was mutated and the mutations were analyzed in enzyme kinetic detail with the transacylation assay. With this approach, we sought to assess the impact of modulated/non-cognate interfaces for the first time in full enzymatic detail. Three available AT:ACP complex structures were used for selecting interface mutations (**Figure 7**). ^[22], [51], [52]^ Eventually two residues, A539 and R850, were selected that are likely involved in the AT–ACP interaction. Overall, nine point mutations were introduced in the AT domain of the KS^0^-AT didomain construct of DEBS3M5; A539S, A539D, A539E, A539F, R850K, R850A, R850E, R850F, R850S. All mutants showed wild type-like properties during preparation: For typical expression yields and SDS-PAGE analyses, see **Table S7 and Figure S9**. SEC showed mainly tetrameric as well as dimeric oligomers like the wild type, and just partly monomeric species (**Figure S10**). High protein quality was confirmed by TSA; i. e., melting temperatures were around 48 °C in storage buffer, which is comparable to the wild type’s melting temperature of 51 °C (**Table S8 and Figure S11**).

**Figure 7:**
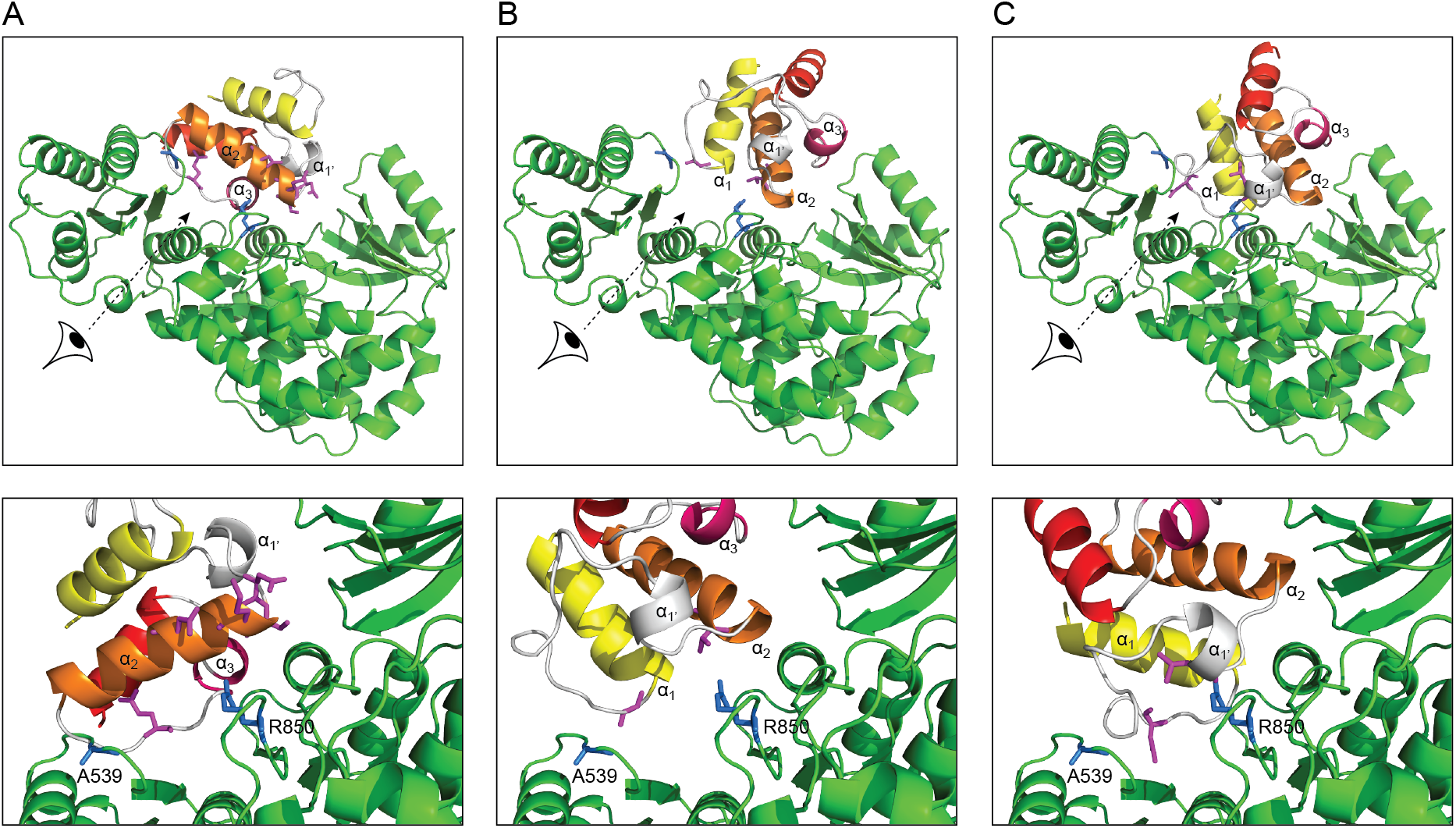
Cartoon representation of the modeled DEBS3M5 AT:ACP interface. DEBS3M5 AT (PDB: 2hg4, shown in green) alignment with the standalone AT from disorazole synthase (PDB: 5zk4) (A), with the standalone AT VinK from vicenistatin synthase (PDB: 5czd) (B) and with MAT from human FAS (C). ^[18], [47], [48]^ Upper panel shows an overview of the AT–ACP interaction of each alignment with perspective eyes for the detailed view showed in the lower panel. Residues A539 and R850 are highlighted in blue. Model DEBS3M5 ACP was aligned on the corresponding ACPs of templates, colored from yellow (N-terminus) to red (C-terminus). Interfacial amino acids suggested by PISA are highlighted in magenta. Despite their different AT:ACP interfaces, all models locate amino acids A539 and R850 (highlighted in blue) in the AT:ACP interfaces.

In an initial screen, the mutants were tested at fixed MMal-CoA and fixed ACP concentrations in transacylation. We also screened the hydrolysis rates at fixed MMal-CoA concentrations. Since this reaction does not involve the interaction of ACP and AT, hydrolysis rates are able to report any impact of the surface mutation on the active site and the binding tunnel. Our initial screen revealed that indeed some mutations affect both transacylation and hydrolysis, although the impact on hydrolysis was comparably lower than on transacylation (**Figure 8**). This data shows that surface point mutations can be invasive to structural and conformational properties that determine the enzymatic reaction. This phenomenon was pronounced for mutations introduced at R850, and not for A539, as depicted in **Figure 8**.

**Figure 8:**
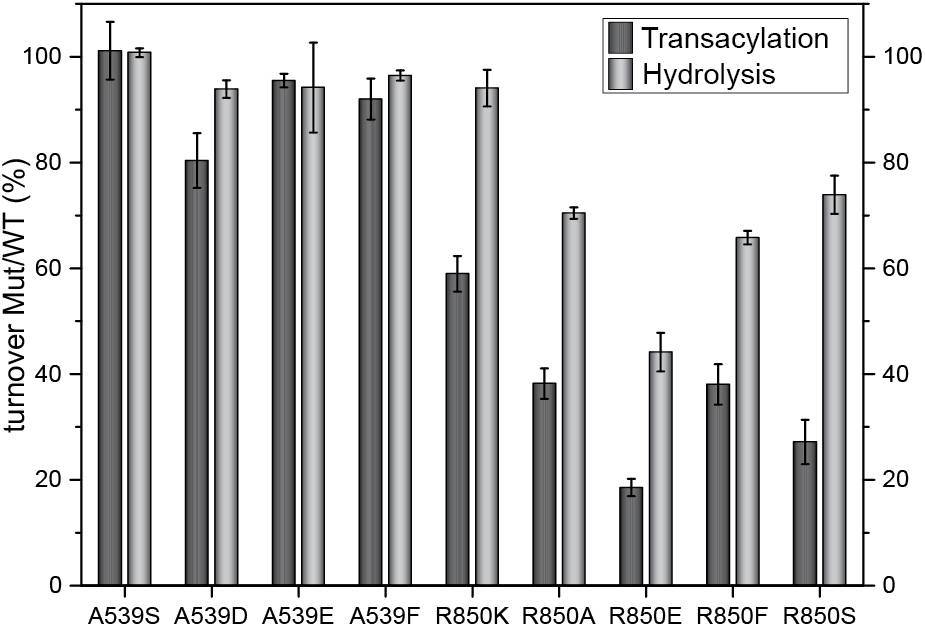
Transacylation (dark grey) and hydrolysis (light grey) screening of the different DEBS3M5 AT:ACP point mutants. The transacylation activity is measured in technical triplicates of one biological replicate; hydrolysis is measured in biological (A539E, R850K, R850E, R850S) or technical triplicates (A539S, A539D, A539F, R850A, R850F). The average activity of each mutant (Mut) is divided by wild type’s (WT) activity and is given in %. Error bars indicate the standard deviation of technical or biological triplicates. A539 mutants seem to have no influence on transacylation and hydrolysis, whereas R850 mutants decrease both transacylation and hydrolysis rates significantly.

Based on the initial screening, four mutations were chosen for full kinetic characterization (in biological triplicates); i. e., A539E, which shows nearly no effect in screening, and the mutations R850K, R850E and R850S, which show moderate to strong effects, preserving, inverting and neutralizing the charge of the mutated residue. For protein quality control of biological replicates, see **Table S8 and Figures S9, S10 and S11**. The ACP concentrations for the enzymatic analysis of mutant R850E lie within a non-ideal range of 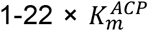 due to a low 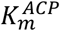 value, and the kinetic transacylation parameters for this mutant could just be determined at lower confidence.

Transacylation and hydrolysis turnover rates well reflect the initial screening (**Table 3 and Figure 9,**for hydrolysis and transacylation titration curves, see **Figure S12**and **S13**). Mutant A539E showed comparable transacylation rates to the wild type protein (82 % of wild type), whereas R850 mutants were significantly compromised in transacylation activity (R850K with 57 % of wild type, R850E 5.5 % and R850S 23 %). Hydrolysis rates remained unaltered by the mutations A539E, R850K and R850S, but dropped to 68 % of wild type rates for mutation R850E. We note (again) that the transacylation rate always includes the hydrolysis rate to some extent.

**Table 3.** Transacylation and hydrolysis kinetic parameters determined for DEBS3AT5 WT and mutants with MMal-CoA. For hydrolysis and transacylation titration curves of all mutants, see **Figure S11 and S13**. Overall, transacylation and hydrolysis kinetic parameters are in good accordance and the catalytic efficiencies give similar values for each mutant, as it should be.

| Transacylation |  |  |  |  |  | Hydrolysis |  |  |
| --- | --- | --- | --- | --- | --- | --- | --- | --- |
| Mutation | $k_{cat}$ ($s^{-1}$) | $K_m$, MMal-CoA ($\mu M$) | $K_m$, ACP ($\mu M$) | $k_{cat}/K_m$, MMal-CoA ($M^{-1}s^{-1}$) | $k_{cat}/K_m$, ACP ($M^{-1}s^{-1}$) | $k_{cat}$ ($s^{-1}$) | $K_m$, MMal-CoA ($\mu M$) | $k_{cat}/K_m$, MMal-CoA ($M^{-1}s^{-1}$) |
| – | $1.25 \pm 7.11 \times 10^{-2}$ | $209 \pm 17.5$ | $72.0 \pm 8.21$ | $6.0 \times 10^3$ | $1.7 \times 10^4$ | $5.31 \times 10^{-2} \pm 9.62 \times 10^{-4}$ | $7.88 \pm 3.98 \times 10^{-1}$ | $6.7 \times 10^3$ |
| A539E | $1.02 \pm 5.19 \times 10^{-2}$ | $164 \pm 13.5$ | $74.3 \pm 7.96$ | $6.2 \times 10^3$ | $1.4 \times 10^4$ | $5.16 \times 10^{-2} \pm 1.15 \times 10^{-3}$ | $9.99 \pm 6.06 \times 10^{-1}$ | $5.2 \times 10^3$ |
| R850K | $7.10 \times 10^{-1} \pm 5.82 \times 10^{-2}$ | $121 \pm 16.3$ | $73.2 \pm 12.7$ | $5.9 \times 10^3$ | $9.7 \times 10^3$ | $5.38 \times 10^{-2} \pm 1.59 \times 10^{-3}$ | $11.2 \pm 8.75 \times 10^{-1}$ | $4.8 \times 10^3$ |
| R850E | $6.82 \times 10^{-2} \pm 3.04 \times 10^{-3}$ | $75.5 \pm 7.65$ | $2.38 \pm 0.408$ | $9.0 \times 10^2$ | $2.9 \times 10^4$ | $3.59 \times 10^{-2} \pm 1.79 \times 10^{-3}$ | $37.7 \pm 5.37$ | $9.5 \times 10^2$ |
| R850S | $2.82 \times 10^{-1} \pm 1.41 \times 10^{-2}$ | $140 \pm 11.7$ | $46.0 \pm 4.55$ | $2.0 \times 10^3$ | $6.1 \times 10^3$ | $5.32 \times 10^{-2} \pm 2.94 \times 10^{-3}$ | $25.9 \pm 3.60$ | $2.1 \times 10^3$ |

**Figure 9:**
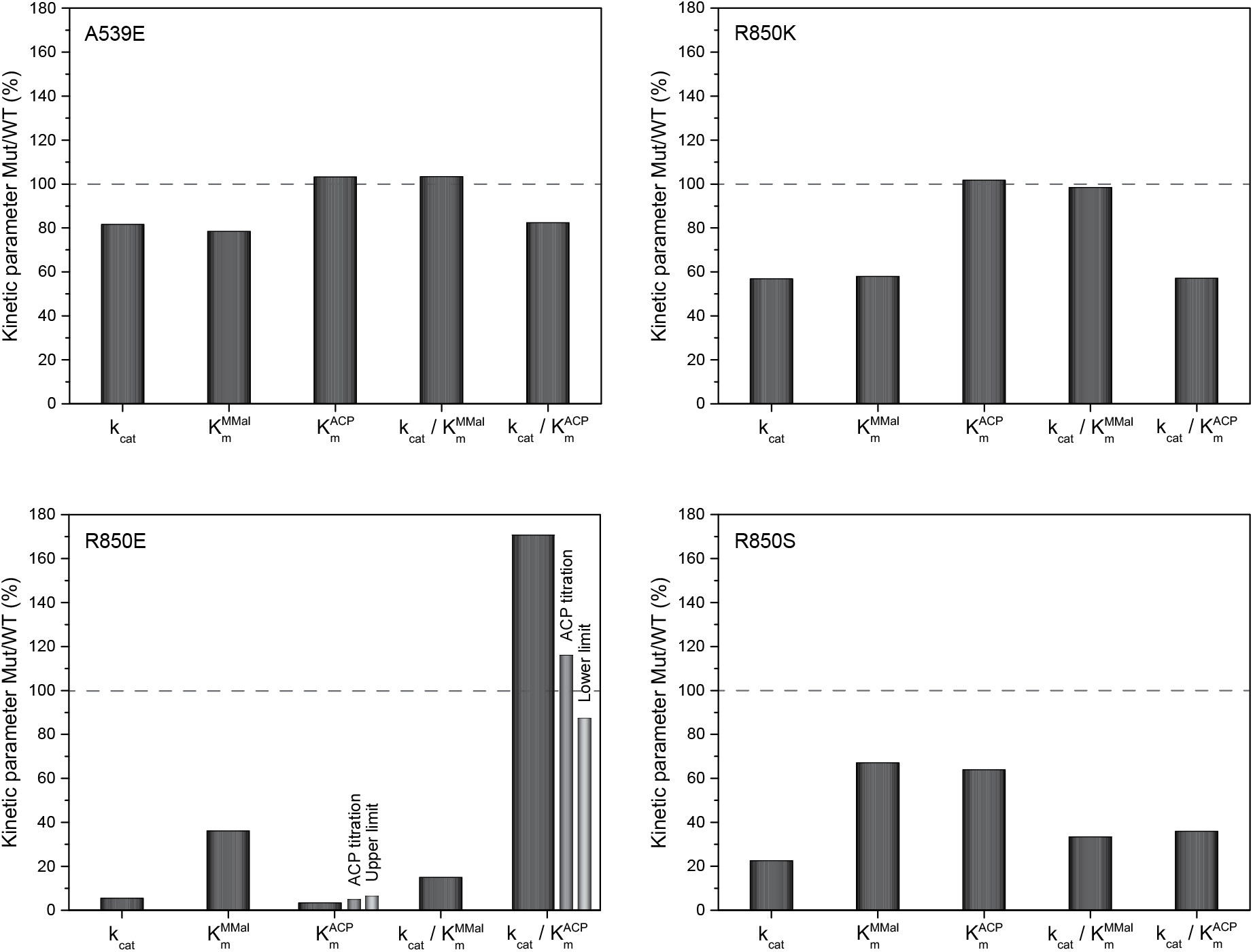
Ratio of kinetic constants determined for transacylation catalyzed by mutants (Mut) A539E, R850K, R850E, and R850S and wild type (WT) kinetic constants in %. R850E shows three ratios of 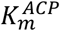 Mut/WT corresponding to the different 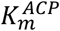 of the mutant: value determined via global fit (ratio shown in dark grey), determined via ACP titration without lowest ACP concentration (ratio shown in grey), and the upper limit of the value (ratio shown in light grey). This gives three corresponding ratios for the catalytic efficiency 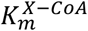.

Overall, the kinetic characterization indicates that the site A539 is rather not involved in the AT–ACP and the AT–MMal-CoA interaction or mutations are not invasive to the AT–ACP interaction. The kinetics of the transacylation reaction is essentially unaltered compared to the wild type, as can also be seen in transition state energies for the ping step and pong step remaining at wild type level (**Figure 10**). In contrast, the mutations of R850 led to larger and more complex effects on both steps. (i) The mutation R850K, preserving the positive charge, has almost no influence on the AT–MMal-CoA interaction, as indicated by the ping step remaining unaltered in catalytic efficiency and transition state energy compared to wild type (**Figure 10**). In contrast, the pong step drops in efficiency, and overall reduces turnover rates for transacylation (57 % of wild type). The 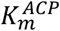 of the pong step is similar to wild type protein, which indicates unchanged AT-X:ACP complex stability. Since the catalytic rate of the pong step (*k_4_*) is decreased, 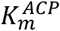 can just remain at wild type level when compensated by an increase of ratio *k_-3_/k_3_*, indicating slower formation (*k_3_*) and/or faster non-productive dissociation of AT-X:ACP (*k_-3_/k_3_*). Overall, mutant R850K specifically affects the pong-step via thermodynamic and kinetic effects. (ii) In the mutant R850S, both ping step and pong step are affected by the amino acid exchange, as indicated by changed catalytic efficiencies and transition state energies compared to wild type, and effects cumulate to a decrease in turnover rates to 23 % compared to wild type. The decrease in the catalytic rate of the ping step (*k_3_*) and the pong step (*k_3_*) account for the increased stabilities of the complexes AT:X-CoA and AT-X:ACP (lowered 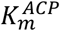 and 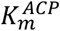 values), but ratios *k_-3_/k_3_* and *k_-3_/k_3_* may contribute here. Overall, the effect of mutation R850S is not constrained to the pong step, but also invasive to AT-X formation; i. e., not showing the specific properties expected of a AT:ACP interface mutation. (iii) The mutant R850E inverses the charge from positive to negative, and we expected severe changes in the kinetics of the transacylation reaction. Indeed, we observed lowest turnover rates within our set of mutants. We note that the kinetic parameters of the pong step could only be collected with lower confidence due to the low 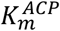. For a reliable estimate on 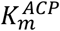, we have therefore extracted data for saturated substrate MMal-CoA concentration titrated with ACP, and receive a 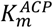 of 3.46 μM and an upper boundary of 4.6 μM (**Figure S14**). Taking this data, we conclude that the pong step remains essentially unaltered in catalytic efficiency and transition state energy. In contrast, the ping step is severely affected by this mutation and responsible for the overall decrease of turnover rates. Since the catalytic rate of the ping step (*k_3_*) is contributing to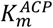, the drop of 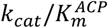 is “just” a direct consequence of the slow ping step. Under this assumption, the kinetics of the pong step would be entirely unaffected in spite of exchanging the positively charged arginine with the negatively charged glutamate. In conclusion, this mutant, designed for a severe impact on the domain-domain interplay during the pong step, mainly (or entirely) takes effect on the initial ping step, which demonstrates the intricate effect surface mutations can have.

**Figure 10:**
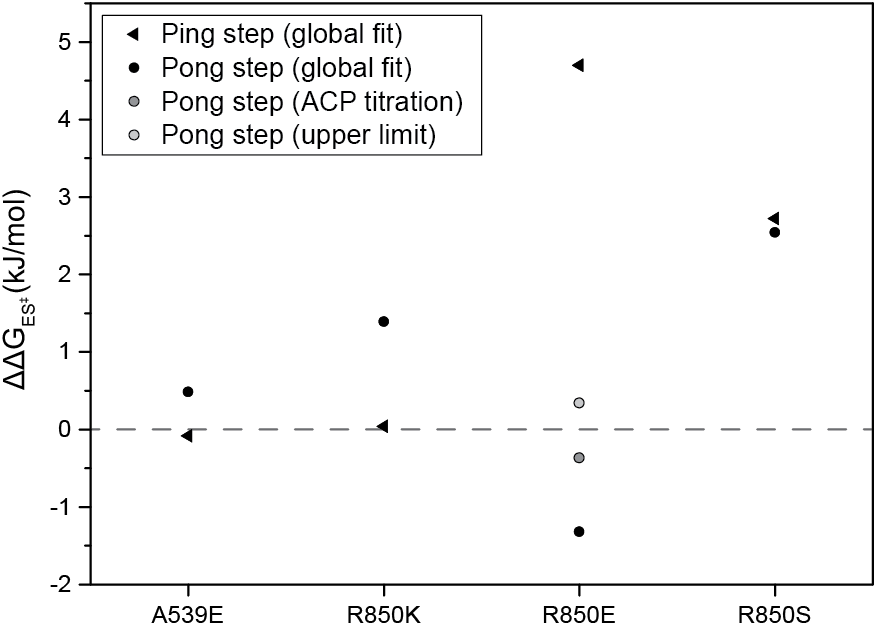
Difference in transition state energy (kJ/mol) between mutants A539E, R850K, R850E, and R850S and wild type determined for the ping step (triangels) and the pong step (dots) of transacylation. Mutant R850E gives three values for 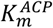: one determined via global fit, one via ACP titration and one upper limit. Differences in transition state energy to wild type are shown in dark grey, grey, and light grey, respectively. 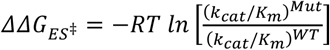 with the gas constant R, T=298 K, and the ratio of catalytic efficiencies between mutant (Mut) and wild type (WT).

### Implications of Interface Mutation Study on PKS Engineering

Electrostatic networks have previously been shown to dominate interactions of ACP with catalytic domains. ^[31], [49]^ Accordingly, we assumed that R850 with its cationic guanidinium headgroup is likely responsible for a such a key electrostatic interaction at the AT:ACP interface. Further support for R850 being involved in AT–ACP interaction was received from X-ray data in which residue R850 is poorly resolved in electron density in the KS-AT didomain structure, ^[28]^ indicating that R850 is not part of an electrostatic network of AT, but involved in AT–ACP interaction (**Figure S15**). Our kinetic data reflect the importance of the positive charge, by the rather moderate effect of R850K mutation on the overall rate of transacylation, compared to mutations R850S and R850E erasing and inverting the charge at this amino acid position, respectively. However, with help of the quantitative data, we likewise revealed that the mutations are not constrained to the AT:ACP interplay, but impose thermodynamic and kinetic changes on the entire reaction sequence (ping step and pong step). Engineering domain-domain interplay can be useful when non-cognate domains are to be integrated in chimeric PKSs. Beyond the boundaries of the DEBS3M5 AT that has served as a model system in this study, the intricate impact of mutations on the catalytic properties suggests that simple engineering strategies based on local effects of amino acid exchanges are prone to fail. For example, engineering interfaces for stabilizing enzyme-substrate complexes (low *k_m_* values) will not necessarily lead to high turnover rates and catalytic efficiencies. The design of chimeric PKSs that perform product synthesis at high efficiency will require deep quantitate analysis and intricate tuning of the enzyme’s kinetic network, ideally complemented by computational methods, as recently demonstrated with FAS. ^[53]^

## Conclusion

FASs and PKSs are evolutionary related enzymes. Their AT domains are key players in biosynthesis and responsible for loading acyl substrates onto the ACP domains of the multidomain complexes. Our findings propose that iterative PKSs and FASs generally feature AT domains with high transacylation rates of 15 to several thousand molecules per second, and show a broad tolerance toward non-natural substrates. In contrast, the evolution of the intricate modular PKSs has led to a significant decrease in turnover rates and substrate tolerance of the AT domains. Today, they act as highly specific selectivity filter during biosynthesis, just transferring 0.7-7 molecules per second. This specificity is in general not assured by hydrolytic proofreading.

Rational engineering of PKSs is a long-standing goal in the field of combinatorial biosynthesis, and envisions generation of novel products in a predictable and controlled manner. The transacylation reaction is an essential setscrew in order to produce new products, and its fine-tuning is possible by the wide range of kinetic properties of AT domains, as revealed in this study. The AT domains of modular PKSs can be used for the loading of specific extender substrates and site-specific modulation of polyketides. In contrast, the substrate promiscuous ATs of iterative systems allow condensing a broader range of extender substrates with product output spectra essentially defined by the substrate specificity of KS and the processing domains. The promiscuous MAT domain of the mammalian FAS seems ideal for such approaches. Since this protein is also evolutionary close to a FAS/PKS common ancestor, it may further represent a well-suited starting point for engineering specificity for new and non-native substrates. ^[54]^

Finally, we show that mutations at the interface between AT and ACP can lead to complex kinetic outputs, particularly when affecting the residues involved in the electrostatic network that guides and tunes enzyme–substrate interactions. Replacing an arginine by lysine, serine or glutamate (R850K, R850S, R850E) led to a severe drop in enzyme-substrate-complex stability, catalytic efficiencies and turnover rates. The impact of mutations can be explained by changing conformational properties that include those of active site and/or binding tunnel distant to the point mutation. The broadly changing kinetic parameters upon single mutations point towards a plasticity in the interaction of AT with acyl-CoA and ACP, as recently suggested for the AT–ACP interaction in the *E. coli* type II FAS (FabD and AcpP). These insights will improve rational PKS engineering approaches.

## Acknowledgements

This project was supported by the LOEWE program (Landes-Offensive zur Entwicklung wissenschaftlich-ökonomischer Exzellenz) of the state of Hesse and was conducted within the framework of the MegaSyn Research Cluster. This project was further supported by an individual PhD scholarship from the Studienstiftung des deutschen Volkes and from the Polytechnische Gesellschaft.

We thank C. S. Heil for carefully reading and discussing the manuscript. We thank G. Grammbitter for HPLC-MS measurements and assistance with analysis of the data. We thank A. Stegemann for discussion of derivation of kinetic parameters. We thank M. Schwalm for assistance with depiction of electron density data. We thank A. Rittner for providing the plasmids pAR001, pAR090, pAR225, pAR331 and pAR333. We thank A. Fahim for performing some SEC und TSA measurements under supervision of F.S.

## Author contributions

F.S. performed all experiments, conceived the project, analyzed data and wrote the manuscript. M.G. designed research, analyzed data and wrote the manuscript.

## Material and Methods

### Plasmid Construction

Coding sequence for FAS and PKS constructs were cloned from genomic DNA into pET22b expression vector, performing ligation independent cloning with the In-Fusion HD Cloning Kit (Clonetech). Single point mutations were introduced in the DEBS3M5 KS^0^-AT wild type gene. The sequence of all plasmids was confirmed by sequencing (Seqlab). ACP sequences were partially codon optimized for expression in *E. coli* and synthesized by Thermo Fisher. All cloning was done using *E. coli* Stellar competent cells.

### Protein Isolation

All constructs were expressed in *E. coli* BL21gold (DE3) cells. All ACPs were co-expressed with *Bacillus subtilis* Sfp. After induction with 0.25 mM IPTG, expression was carried out at 20 °C and 130-180 rpm overnight. Transferases were purified by tandem affinity chromatography (Ni-NTA and Strep-Tactin), ACPs were purified by Ni-affinity chromatography and size exclusion chromatography (SEC). ACP from different expression cultures was pooled after SEC. Elution fractions were analyzed by SDS-PAGE. Quality of transferases was analyzed by HPLC-SEC and thermal shift assay (TSA). Activation of ACPs was controlled by mass spectrometric analysis.

### AT Activity Assay

The AT activity assay was adapted from references. ^[13], [16], [24]^ For all proteins and substrates, fresh aliquots were used. For each system the same batch of substrates (ACP and acyl-CoAs) were used. For better comparison, for all point mutants the same batch of substrates was used. NADH fluorescence was detected in a 96-well format at fixed AT and ACP concentrations. For hydrolysis and for each ACP curve in transacylation measurements, acyl-CoA substrate concentration was varied to calculate the (apparent)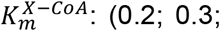. 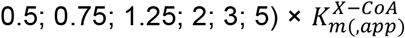 Fluorescence units were converted into concentrations via NADH calibration (**Figure S16**).

