## Supporting Information for "Comparative Study on Acyl Transferases in Fatty Acid and Polyketide Synthases"

**Table S1.** Average expression yields of AT domains after tandem-affinity chromatography. Yields determined from three independent expressions.

| Protein | Yield (mg )per 1 L expression culture |
|---|---|
| DEBS3 (KS$^0$-AT)5 | 14.0 |
| PikAIII (KS$^0$-AT)5 | 13.8 |
| RAPS3 (KS$^0$-AT)14 | 3.47 |
| Pks7 KS$^0$-AT | 21.5 |

**Table S2.** TSA of AT and ACP domains. Average of biological triplicates (ATs) and technical triplicates (ACPs).

| Protein | $T_m$ (°C) | | | |
|---|---|---|---|---|
| | AT | | ACP | |
| | Storage Buffer | Assay Buffer | Storage Buffer | Assay Buffer |
| DEBS3M5 | 51 | 42 | 64 | 65 |
| PikAIIIM5 | 49 | 42 | 59 | 56 |
| RAPS3M14 | 45 | 36 | 57 | 50 |
| Pks7 | 49 | 41 | 62 | 58 |

**Table S3.** Average expression yields of non-codon optimized and codon optimized (COP) *holo*-ACP domains after SEC. Yields determined from four to nine independent expressions.

| Protein | Yield (mg) per 1 L expression culture |
|---|---|
| DEBS3 ACP5 | 23 |
| DEBS3 ACP5 COP | 87 |
| PikAIII ACP5 | 4.5 |
| PikAIII ACP5 COP | 51 |
| RAPS3 ACP14 | 31 |
| Pks7 ACP | 1.0 |
| Pks7 ACP COP | 30 |

**Table S4.** MS data of ACPs. m/z expected (exo) and found experimentally of *holo*- and *apo*-ACP with and without N-terminal Methionine (Met). All ACPs are fully activated (see Figure S5).

| Protein | m/z (-Met) | | | | m/z (+Met) | | | | State of Charge |
|---|---|---|---|---|---|---|---|---|---|
| | *apo* | | *holo* | | *apo* | | *holo* | | |
| | exp | found | exp | found | exp | found | exp | found | |
| DEBS3ACP5, WT | 836.21 | – | 855.15 | 855.16 | 843.49 | – | 862.43 | – | 18 |
| DEBS3ACP5, Mut | 836.21 | 836.19 | 855.15 | 855.14 | 843.49 | – | 862.43 | – | 18 |
| PikAIIIACP5 | 822.92 | – | 840.82 | 840.81 | 829.20 | – | 848.15 | – | 18 |
| RAPS3ACP14 | 767.88 | – | 786.78 | 786.78 | 775.16 | – | 794.06 | – | 18 |
| Pks7 ACP | 891.29 | – | 910.18 | 910.20 | 898.57 | – | 917.46 | – | 18 |

**Table S5.** Transacylation and hydrolysis rates of initial substrate screening (50 µM X-CoA and ACP). Transacylation measured in biological triplicates. Hydrolysis measured in technical triplicates of each biological triplicate. Abbreviation: Ac: acetyl, Mal: malonyl, MMal: methylmalonyl.

| Protein | X-CoA | Transacylation $k_{0,app}$ ($s^{-1}$) | Hydrolysis $k_0$ ($s^{-1}$) |
|---|---|---|---|
| DEBS3AT5 | MMal | $1.92 \times 10^{-1}$ | $4.24 \times 10^{-2}$ |
| PikAIIIAT5 | Mal | $8.86 \times 10^{-2}$ | $4.98 \times 10^{-2}$ |
| PikAIIIAT5 | MMal | $8.60 \times 10^{-1}$ | $1.54 \times 10^{-2}$ |
| RAPS3AT14 | Mal | $2.05 \times 10^{-1}$ | $6.63 \times 10^{-2}$ |
| mMAT [a] | Ac | 13.7 | $9.3 \times 10^{-3}$ |
| mMAT [a] | Mal | 15.6 | $9.8 \times 10^{-3}$ |
| mMAT [a] | MMal | 4.0 | $6.0 \times 10^{-3}$ |
| Pks7 AT | Ac | $2.81 \times 10^{-1}$ | $1.60 \times 10^{-3}$ |
| Pks7 AT | Mal | 2.79 | $7.05 \times 10^{-2}$ |
| Pks7 AT | MMal | 1.69 | $5.58 \times 10^{-2}$ |

[a] mMAT from previous study, referring to $k_{cat}$ for hydrolysis (saturated X-CoA) and $k_{cat,app}$ for transacylation (saturated X-CoA, 60 µM ACP). [1]

**Table S6.** Kinetic parameters determined for AT-mediated hydrolysis. Turnover measured in technical triplicates of biological triplicates (DEBS3AT5), technical triplicates of one replicate (PikAIIIAT5, RAPS3,AT14), or biological triplicates (Pks7 with Mal and MMal).

| Protein<br>X-CoA | $k_{cat}$ ($s^{-1}$) [a] | $K_{m,\ X\text{-}CoA}$ (μM) [b] | $k_{cat}/K_{m,\ X\text{-}CoA}$ ($M^{-1}s^{-1}$) [c] |
|---|---|---|---|
| DEBS3AT5<br>MMal | $5.31 \times 10^{-2} \pm 9.62 \times 10^{-4}$ | 8.85 | $6.0 \times 10^{3}$ |
| PikAIIIAT5<br>MMal | $1.94 \times 10^{-2} \pm 5.03 \times 10^{-4}$ | $2.49 \times 10^{-1}$ | $7.8 \times 10^{4}$ |
| RAPS3AT14<br>Mal | $6.51 \times 10^{-2} \pm 1.89 \times 10^{-3}$ | 1.36 | $4.8 \times 10^{4}$ |
| Pks7 AT<br>Mal | $9.93 \times 10^{-2} \pm 3.17 \times 10^{-3}$ | $3.97 \times 10^{-2}$ | $2.5 \times 10^{6}$ |
| Pks7 AT<br>MMal | $7.34 \times 10^{-2} \pm 1.51 \times 10^{-3}$ | $7.57 \times 10^{-2}$ | $9.7 \times 10^{5}$ |
| mMAT<br>Mal [d] | $(9.8 \pm 1.7) \times 10^{-3}$ | $7.0 \times 10^{-4}$ | $1.4 \times 10^{7}$ |
| mMAT<br>Ac [d] | $9.3 \times 10^{-3} \pm 1.2 \times 10^{-4}$ | $1.1 \times 10^{-3}$ | $8.3 \times 10^{6}$ |

[a] Value experimentally determined in hydrolysis measurement. [b] Value calculated from experimentally determined hydrolysis and transacylation values. [c] Value experimentally determined in transacylation measurement. [d] mMAT data from previous study. [1]

**Table S7.** Average expression yields of DEBS3 (KS$^0$-AT)5 mutants after tandem-affinity chromatography. Data from three independent protein expressions, except proteins marked with * are single expressions.

| Protein | Yield (mg) per 1 L expression culture |
|---|---|
| A539S* | 10.2 |
| A539D* | 15.4 |
| A539E | 9.67 |
| A539F* | 1.70 |
| R850K | 14.0 |
| R850A* | 5.71 |
| R850E | 7.73 |
| R850F* | 2.36 |
| R850S | 9.62 |

**Table S8.** TSA of DEBS3 (KS$^0$-AT)5 mutants. Average of technical/biological triplicates. DEBS3M5 ACP COP was used in the mutagenesis study.

| Mutation | $T_m$ (°C) | | |
|---|---|---|---|
| | AT | ACP | |
| | Storage Buffer | Storage Buffer | Assay Buffer |
| – | 51 | 63 | 62 |
| A539S | 48 | – | – |
| A539D | 48 | – | – |
| A539E | 48 | – | – |
| A539F | 48 | – | – |
| R850K | 48 | – | – |
| R850A | 48 | – | – |
| R850E | 47 | – | – |
| R850F | 48 | – | – |
| R850S | 48 | – | – |

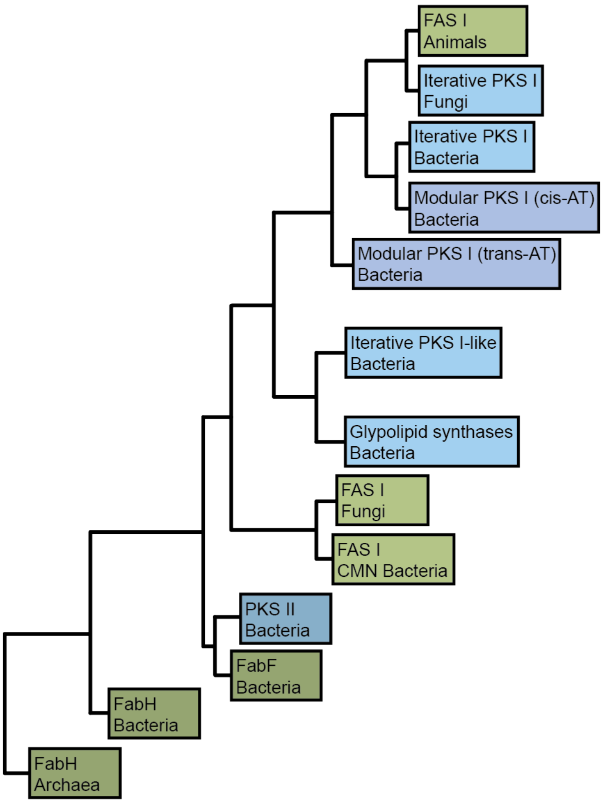

**Figure S1: Schematic representation of the phylogenetic relationship between FASs and PKSs based on the alignment of KS domains.** The type II FAS and PKS systems have predated the multidomain and multimodular type I systems. Type I systems shown in light, type II systems shown in dark colors. FAS systems shown in green, PKS systems shown in blue. Adapted from references. [2], [3]

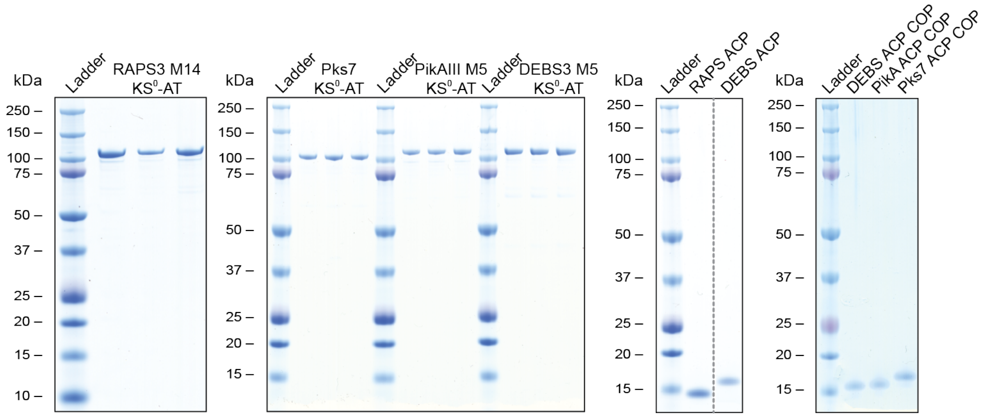

**Figure S2: Analytical SDS-PAGEs of ACP domains and $KS^0$-AT constructs expressed in biological triplicates.** All AT constructs are highly pure after tandem affinity chromatography. Protein bands migrate at expected masses: RAPS3M14 $KS^0$-AT – 101 kDa, Pks7 $KS^0$-AT – 98.6 kDa, PikAIIIM5 $KS^0$-AT – 101 kDa, DEBS3M5 $KS^0$-AT – 99.9 kDa. All pooled ACPs are highly pure after SEC. Masses are as expected: RAPS3M5 ACP – 14.1 kDa, DEBS3M5 ACP – 15.4 kDa, PikAIIM5 ACP – 15.2 kDa, Pks7 ACP – 16.4 kDa.

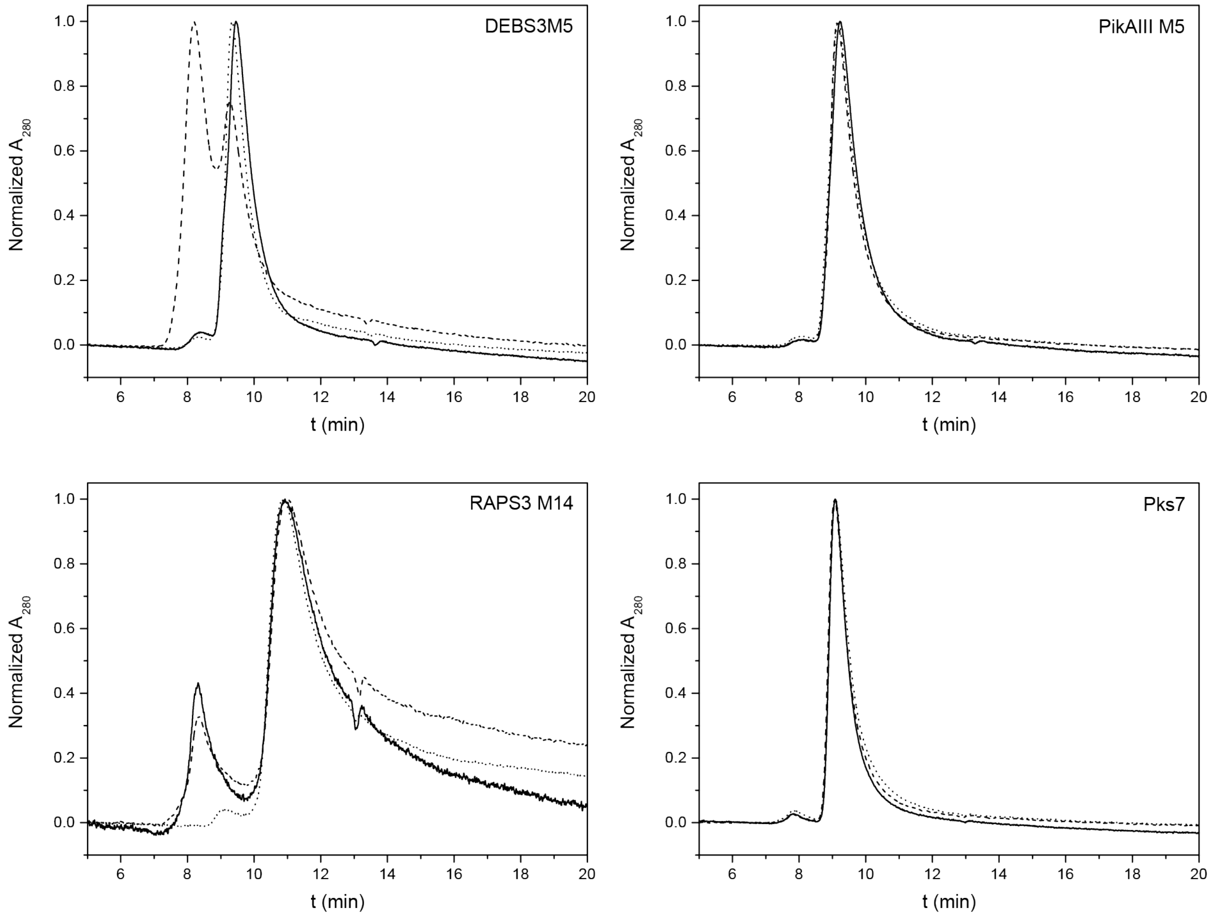

**Figure S3: Normalized size exclusion chromatograms of KS$^0$-AT domains.** Each curve (solid, dotted, dashed) corresponds to one biological replicate. DEBS3M5 KS$^0$-AT shows dimeric (9.4 min) and tetrameric (8.2 min) oligomers. PikAIIIM5 KS$^0$-AT forms solely dimers (9.2 min). RAPS3M14 KS$^0$-AT shows monomeric (11 min) and tetrameric (8.3 min) species. Pks7 KS$^0$-AT forms solely dimers (9.1 min).

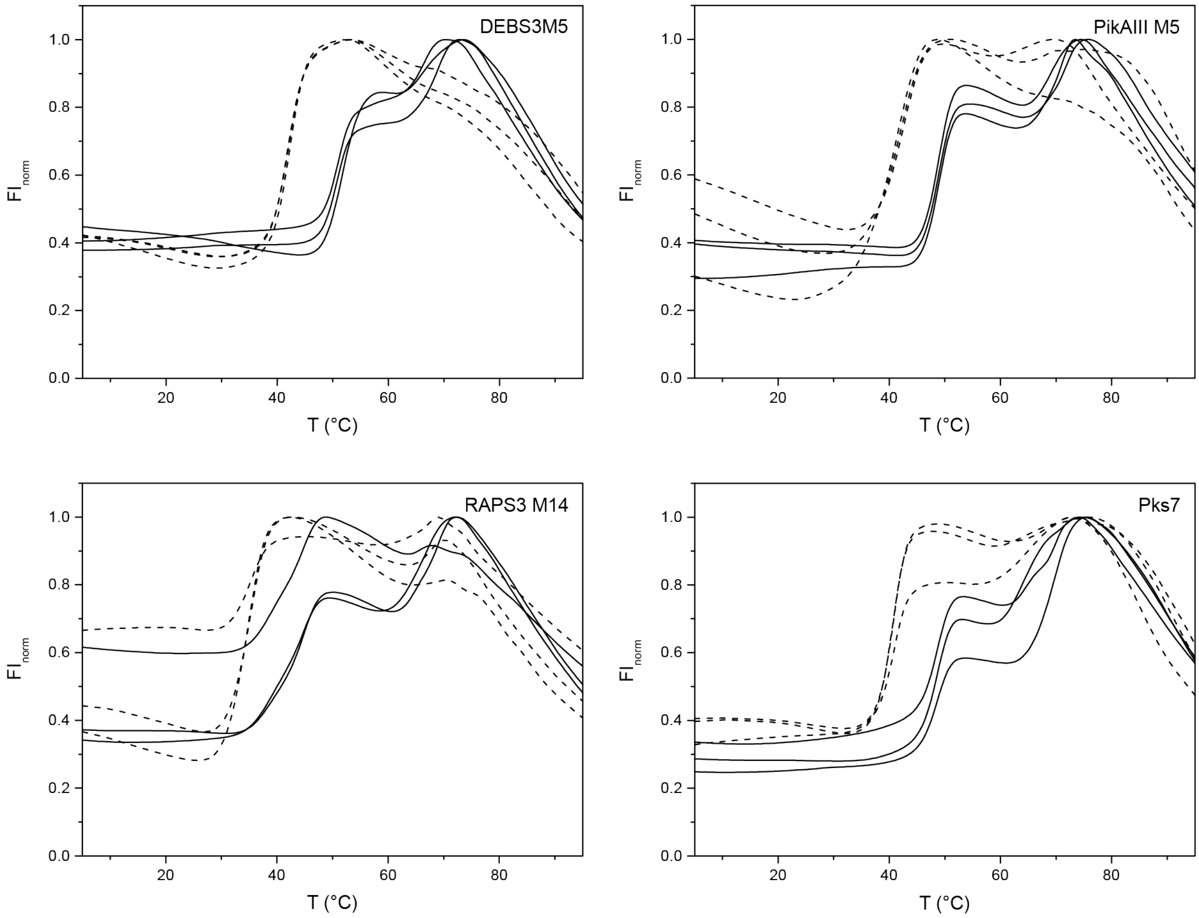

**Figure S4: Normalized TSA melting curves of $KS^0$-AT domains.** Solid and dashed line display data collected in storage and assay buffer, respectively. Each curve (solid, dashed) corresponds to one biological replicate.

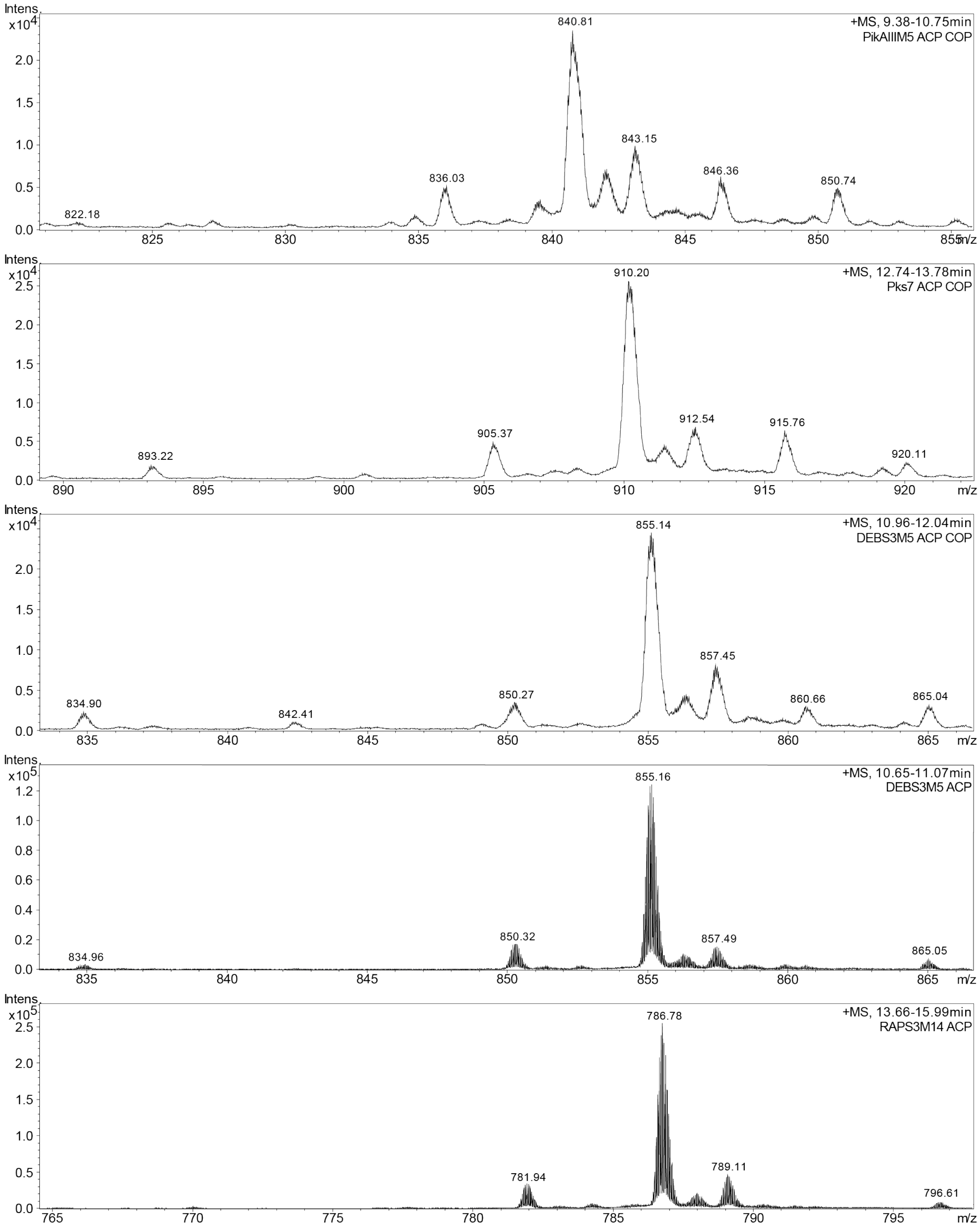

**Figure S5: Mass spectrometric analysis of ACP domains after SEC.** Shown are the m/z-values of the average protein masses of the $18^+$ charge states ($MS^1$). In none of the samples, m/z-values of the *apo* species were recorded. Main m/z-values correspond to the *holo* form. Proteins do not include the N-terminal methionine. COP stands for codon optimized. **PikAIIIM5 ACP COP**, z = 18: m/z = 840.81 corresponds to *holo*-ACP without N-terminal methionine. m/z = 843.15 corresponds to acetylation (Δm = 42 Da), m/z = 850.74 corresponds to α-N-gluconozylation (Δm = 178 Da). m/z = 846.36 is an unknown modified variant (Δm = 100 Da). m/z = 836.03 corresponds to *holo*-ACP without N-terminal methionine and serine ($\Delta m$ = -86 Da). **Pks7 ACP COP**, z = 18: m/z = 910.20 corresponds to *holo*-ACP without N-terminal methionine. m/z = 912.54 corresponds to acetylation ($\Delta m$ = 42 Da), m/z = 920.11 corresponds to α-N-gluconozylation ($\Delta m$ = 178 Da). m/z = 915.76 is an unknown modified variant ($\Delta m$ = 100 Da). m/z = 905.37 corresponds to *holo*-ACP without N-terminal methionine and serine ($\Delta m$ = -87 Da). **DEBS3M5 ACP COP**, z = 18, used for kinetic analysis of mutants: m/z = 855.14 corresponds to *holo*-ACP without N-terminal methionine. m/z = 857.45 corresponds to acetylation ($\Delta m$ = 42 Da), m/z = 865.04 corresponds to α-N-gluconozylation ($\Delta m$ = 178 Da). m/z = 860.66 is an unknown modified variant ($\Delta m$ = 99 Da). m/z = 850.27 corresponds to *holo*-ACP without N-terminal methionine and serine ($\Delta m$ = -88 Da). **DEBS3M5 ACP**, z = 18, used for kinetic analysis of the wild type: m/z = 855.16 corresponds to *holo*-ACP without N-terminal methionine. m/z = 857.49 corresponds to acetylation ($\Delta m$ = 42 Da), m/z = 865.05 corresponds to α-N-gluconozylation ($\Delta m$ = 178 Da). m/z = 860.66 is an unknown modified variant ($\Delta m$ = 99 Da). m/z = 850.32 corresponds to *holo*-ACP without N-terminal methionine and serine ($\Delta m$ = -87 Da). **RAPS3M14 ACP**, z = 18: m/z = 786.78 corresponds to *holo*-ACP without N-terminal methionine. m/z = 789.11 corresponds to acetylation ($\Delta m$ = 42 Da), m/z = 796.61 corresponds to α-N-gluconozylation ($\Delta m$ = 177 Da). m/z = 781.94 corresponds to *holo*-ACP without N-terminal methionine and serine ($\Delta m$ = -87 Da).

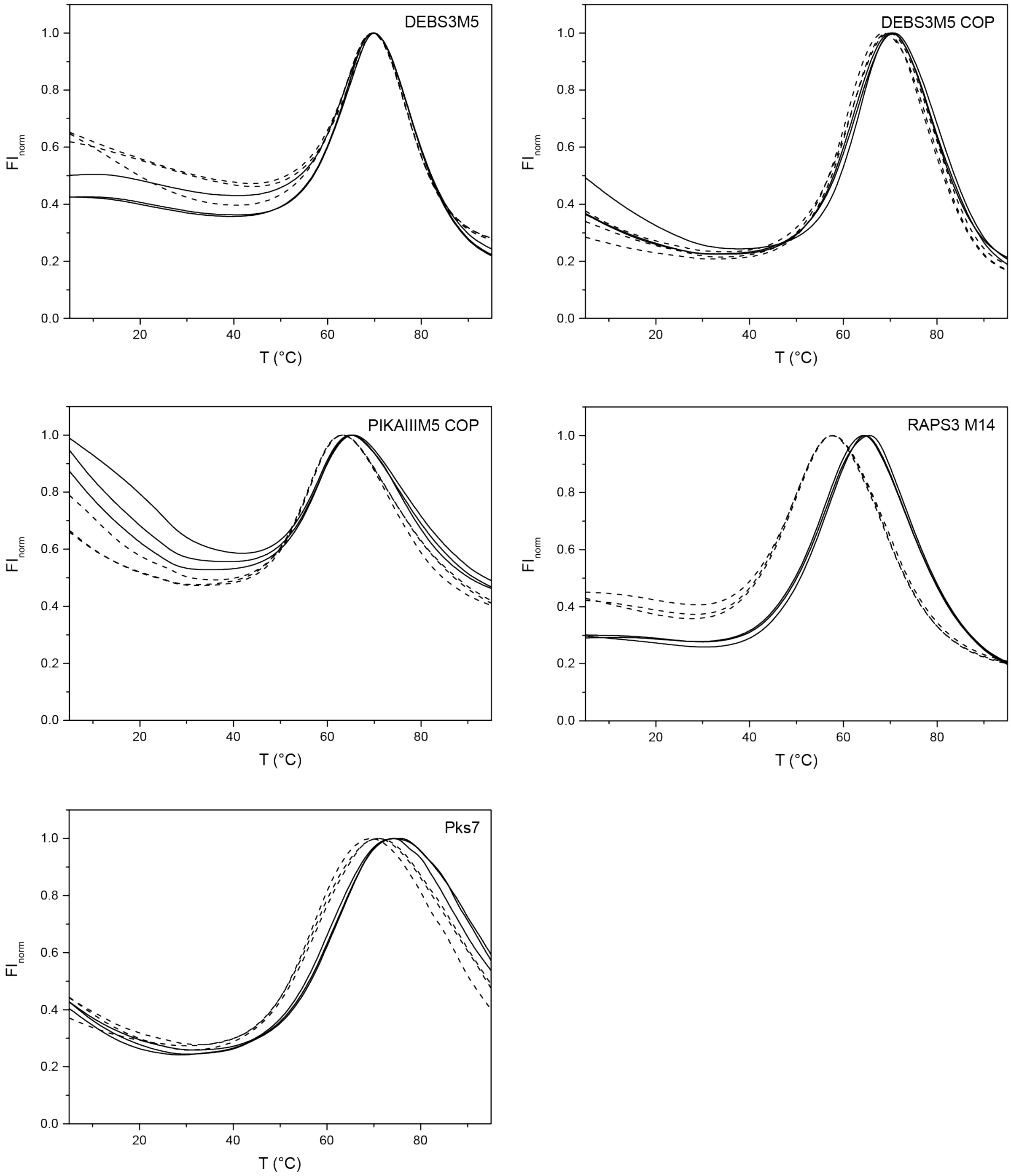

**Figure S6: Normalized TSA melting curves of ACP domains.** Solid and dashed line display data collected in storage and assay buffer, respectively. Each curve (solid, dashed) corresponds to one technical replicate. COP stands for codon optimized.

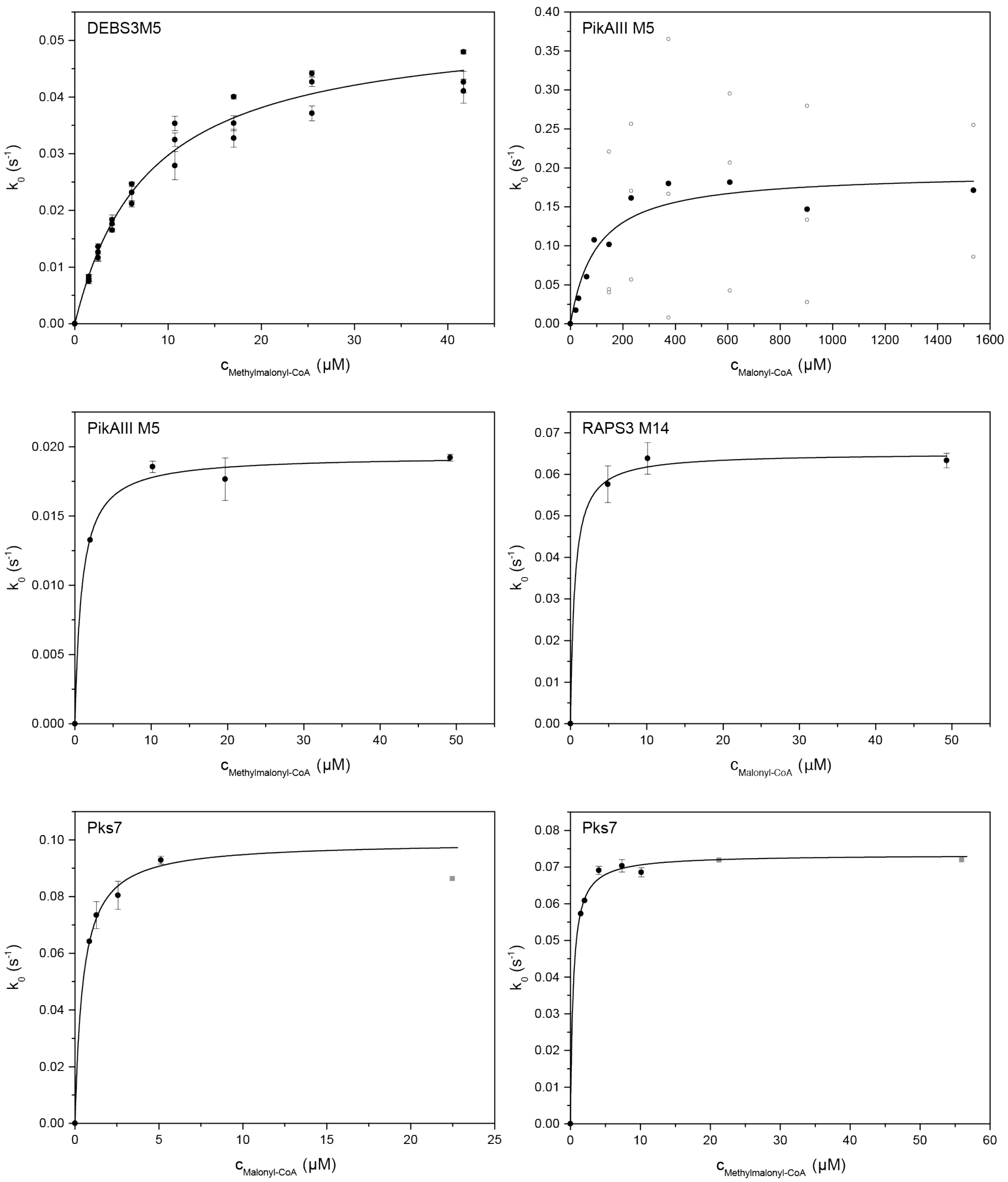

**Figure S7: Titration curves for AT-mediated hydrolysis of X-CoA. DEBS3M5, MMal:** DEBS3M5 KS$^0$-AT measured in technical triplicates of biological triplicates. Black dots with error bars show the average of one biological replicate with standard deviation. Kinetic parameters were determined precisely. Substrate consumption was below 10 %. MMal range was 0.19-5.3 × $K_m^{MMal-CoA}$. **PikAIIIM5, Mal:** PikAIIIM5 KS$^0$-AT measured in biological triplicates. Circles show one biological replicate, black dots show their average. Problems with high Mal concentrations are observed. Only lower limit of kinetic parameters were calculated. Substrate consumption was below 5 %. **PikAIIIM5, MMal:** PikAIIIM5 KS$^0$-AT measured in technical triplicates. Black dots with error bars show the average with standard deviation. The lowest concentration is a one well measurement. Due to low the low substrate concentrations, the maximal velocity and an upper limit of the Michaelis-Menten constant were determined for this system. Substrate consumption was up to 17 %. **RAPS3M14, Mal:** RAPS3M14 $KS^0$-AT measured in technical triplicates. Black dots with error bars show the average with standard deviation. Due to low the low substrate concentration, the maximal velocity and an upper limit of the Michaelis-Menten constant were determined. Substrate consumption was up to 24 %. **Pks7, Mal:** Pks7 $KS^0$-AT measured in biological triplicates. Black dots with error bars show the average with standard deviation. The lowest concentration is a one well measurement. The grey box is a single well measurements with high substrate concentration, which was not used for the hydrolysis fit. It shows that maximal velocity was reached. Due to the low substrate concentrations, the maximal velocity and an upper limit of the Michaelis-Menten constant were determined. Substrate consumption was up to 44 %. Signal increase was still constant. **Pks7, MMal:** Pks7 $KS^0$-AT measured in biological triplicates. Black dots with error bars show the average with standard deviation. The two lowest concentrations are one well measurements. Grey boxes are single well measurements with high substrate concentrations, which were not used for the hydrolysis fit. These values show that maximal velocity was reached. Due to the low substrate concentration, the maximal velocity and an upper limit of the Michaelis-Menten constant were determined. Substrate consumption was up to 34 %. Signal increase was still constant.

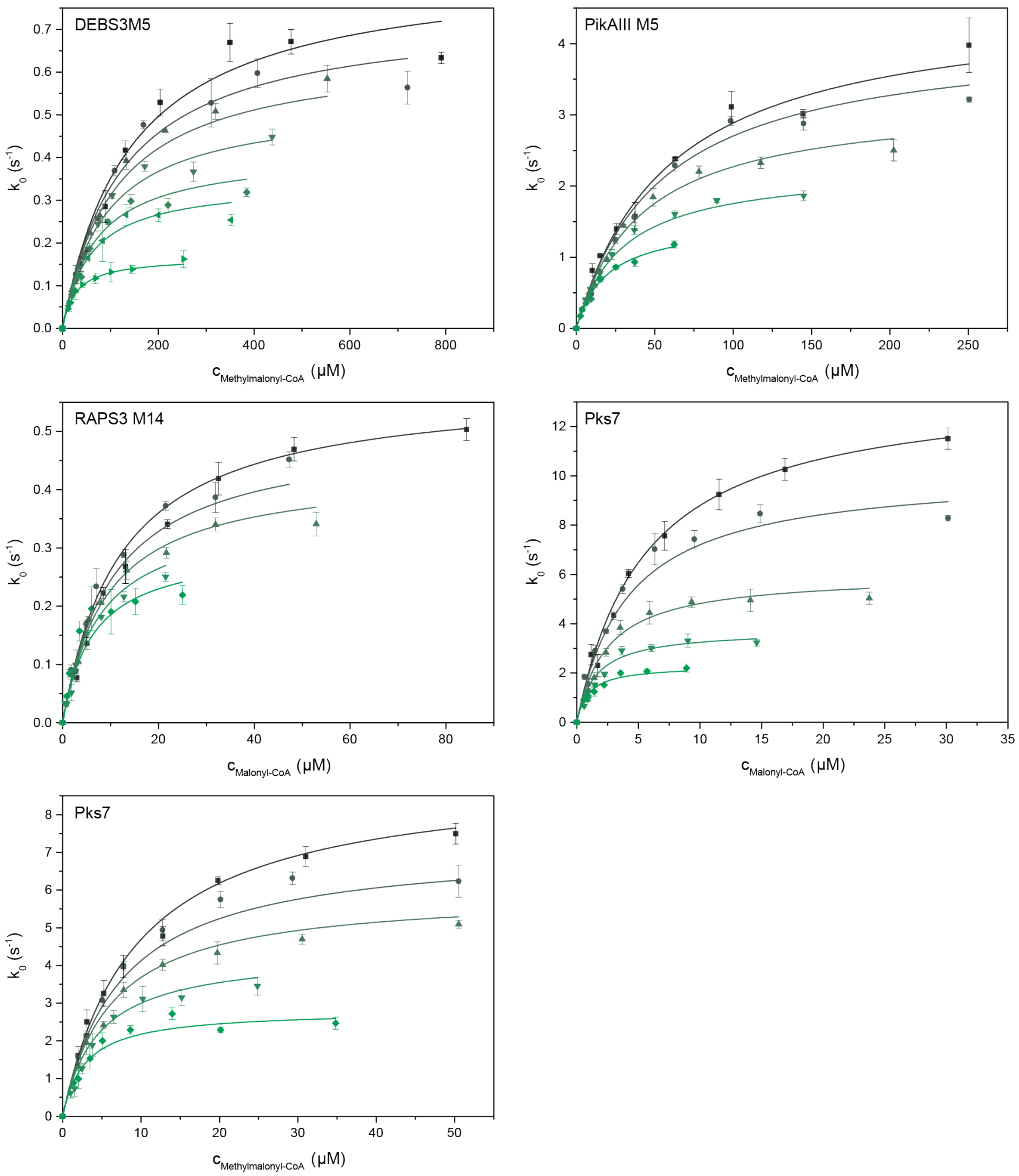

**Figure S8: Global Michaelis-Menten fits of transacylation titration curves with X-CoA of AT domains.** All measured in biological triplicates. Each color and symbol corresponds to one X-CoA titration curve at a fixed ACP concentration. Error bars give the standard deviation of biological triplicates. ACP concentration as follows: DEBS3M5: 11.1 µM, 27.2 µM, 35.4 µM, 53.1 µM, 78.2 µM, 104.9 µM and 152.1 µM, PikAIIIM5: 83.3 µM, 143.9 µM, 249.2 µM, 412.9 µM and 528.4 µM, RAPS3M14: Pks7, Mal: 31.2 µM, 54.5 µM, 98.6 µM, 220.7 µM and 403.6 µM, Pks7, MMal: 80.3 µM, 136.0 µM, 206.3 µM, 273.1 µM and 402.5 µM. ACP concentration ranges as follows: DEBS3M5: 0.15-2.1 × $K_m^{ACP}$, PikAIIIM5: 0.25-1.6 × $K_m^{ACP}$, RAPS3M14: 0.74-4.4 × $K_m^{ACP}$, Pks7, Mal: 0.11-1.4 × $K_m^{ACP}$, Pks7, MMal: 0.16-0.80 × $K_m^{ACP}$. Substrate consumption as follows: DEBS3M5: below 5 %, PikAIIIM5:

below 11 %, RAPS3M14: up to 36 %, but signal increase was still linear, Pks7, Mal: up to 42 %, but signal increase was still linear, Pks7, MMal: up to 18 %. X-CoA concentration ranges as follows: DEBS3M5: 0.16-8.3 × $K_{m,app}^{MMal-CoA}$, PikAIIIM5: 0.12-5.4 × $K_{m,app}^{MMal-CoA}$, RAPS3M5: 0.076-8.7 × $K_{m,app}^{Mal-CoA}$, Pks7: 0.19-9.5 × $K_{m,app}^{Mal-CoA}$ and 0.19-12 × $K_{m,app}^{MMal-CoA}$.

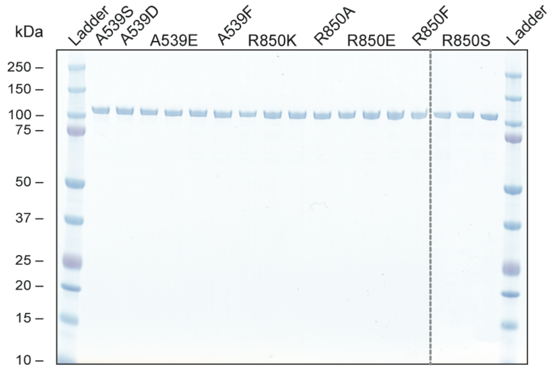

**Figure S9: Analytical SDS-PAGEs of DEBS3M5 KS$^0$-AT mutants.** Mutants A539E, R850K, R850E and R850S expressed in biological triplicates. All AT constructs are highly pure after tandem affinity chromatography. Protein bands migrate at expected masses: A539S – 99.9 kDa, A539D – 99.9 kDa, A539E – 99.9 kDa, A539F – 99.9 kDa, R850K – 99.8 kDa, R850A – 99.8 kDa, R850E – 99.8 kDa, R850F – 99.8 kDa, 850S – 99.8 kDa, R850S – 99.8 kDa.

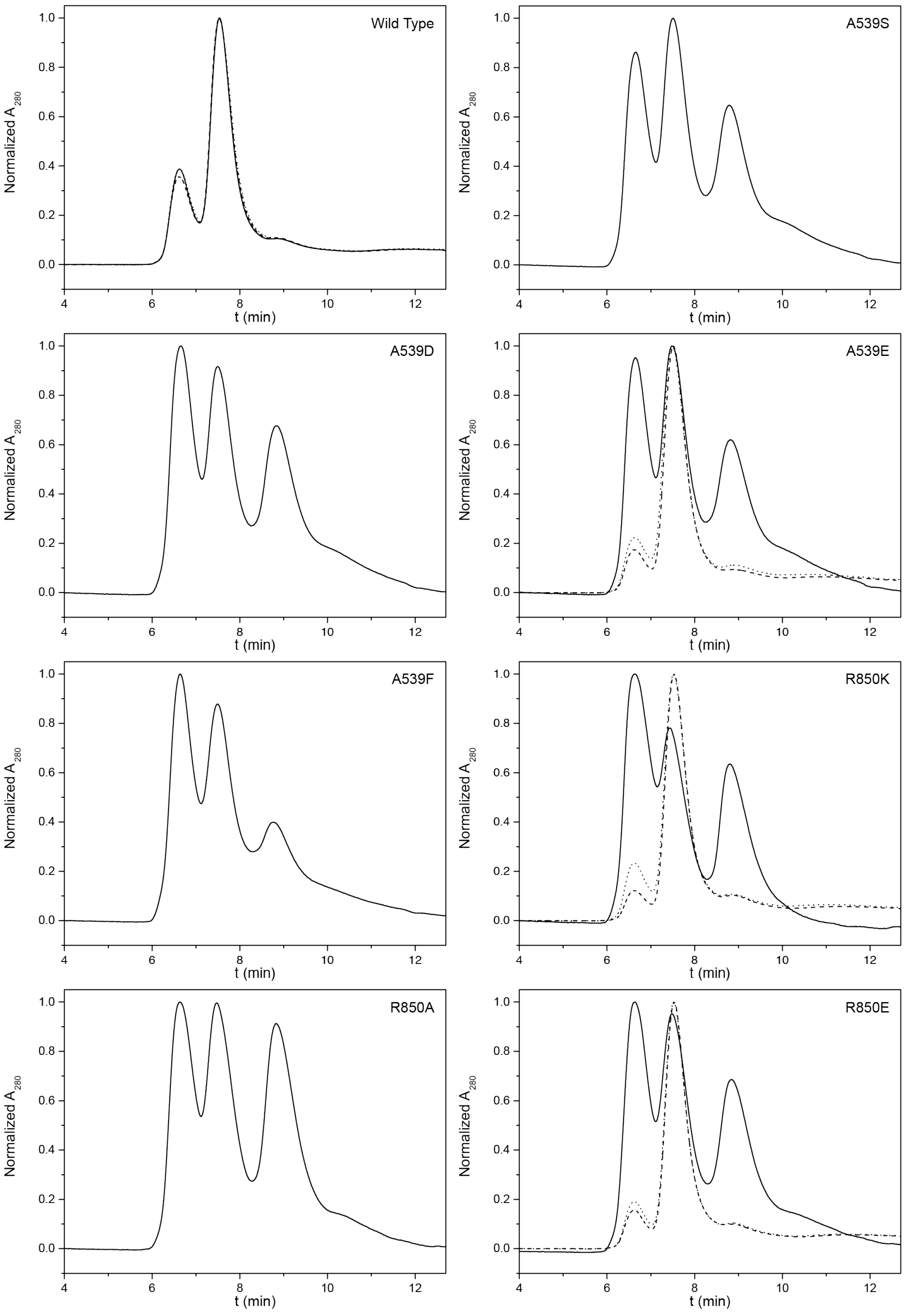
Wild Type
A539S
A539D
A539E
A539F
R850K
R850A
R850E
Normalized A280
t (min)
0.0
0.2
0.4
0.6
0.8
1.0
4
6
8
10
12

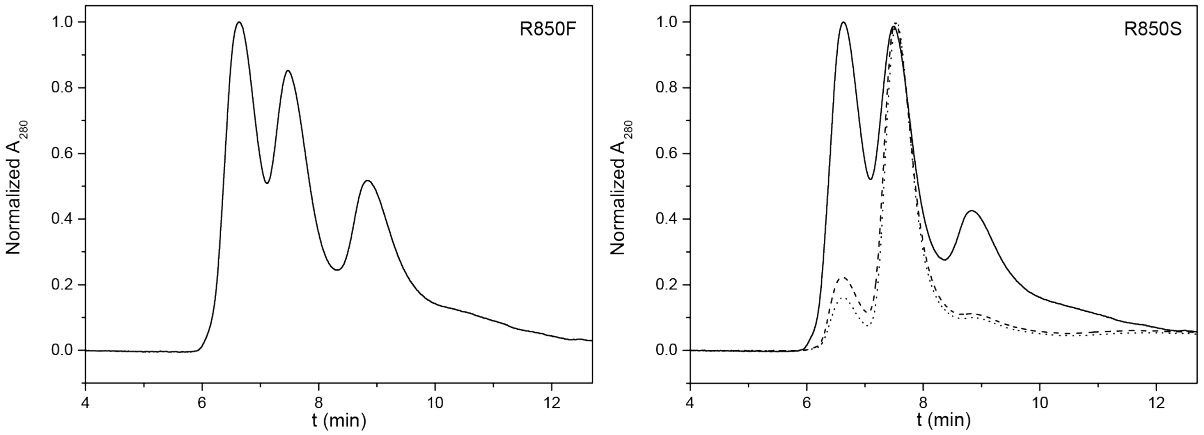

**Figure S10: Normalized size exclusion chromatograms of DEBS3M5 $KS^0$-AT mutants.** Each curve (solid, dotted, dashed) corresponds to one biological replicate. All proteins show monomeric (8.8 min), dimeric (7.5 min) and tetrameric (6.6 min) species.

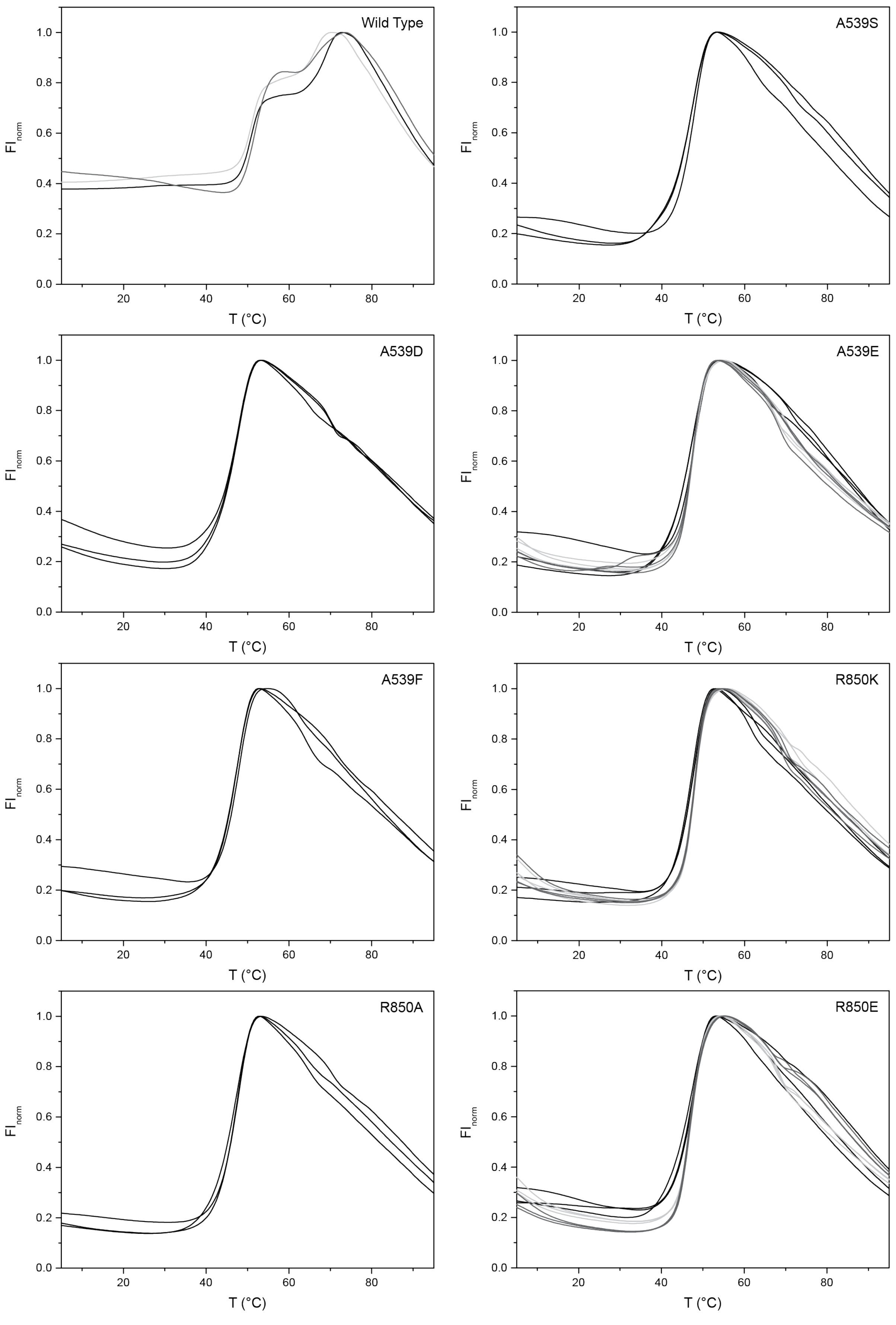
Wild Type
A539S
A539D
A539E
A539F
R850K
R850A
R850E
FInorm
T (°C)
0.0
0.2
0.4
0.6
0.8
1.0
20
40
60
80

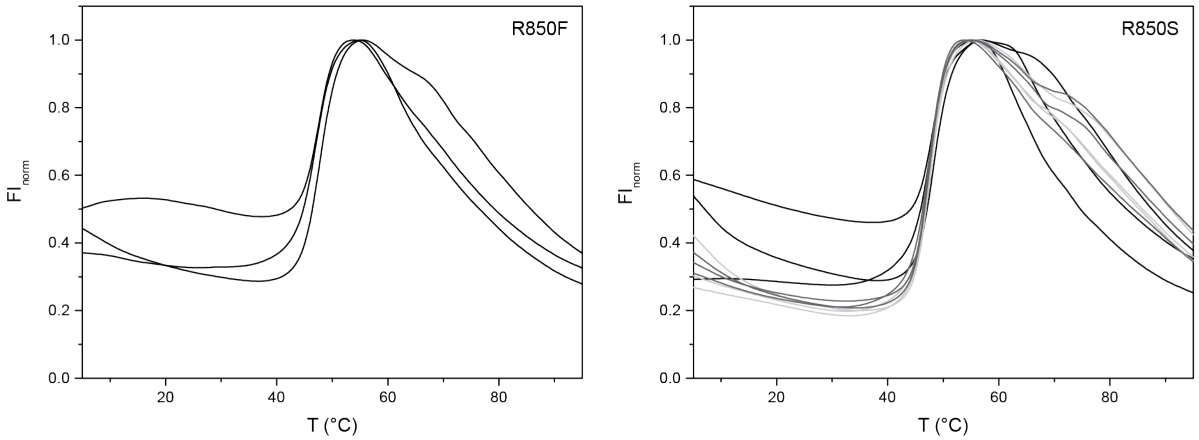

**Figure S11: Normalized TSA melting curves of DEBS3M5 $KS^0$-AT mutants.** Each color (black, dark and light grey) corresponds to one biological replicate. All biological replicates of mutants measured in technical triplicates in assay buffer.

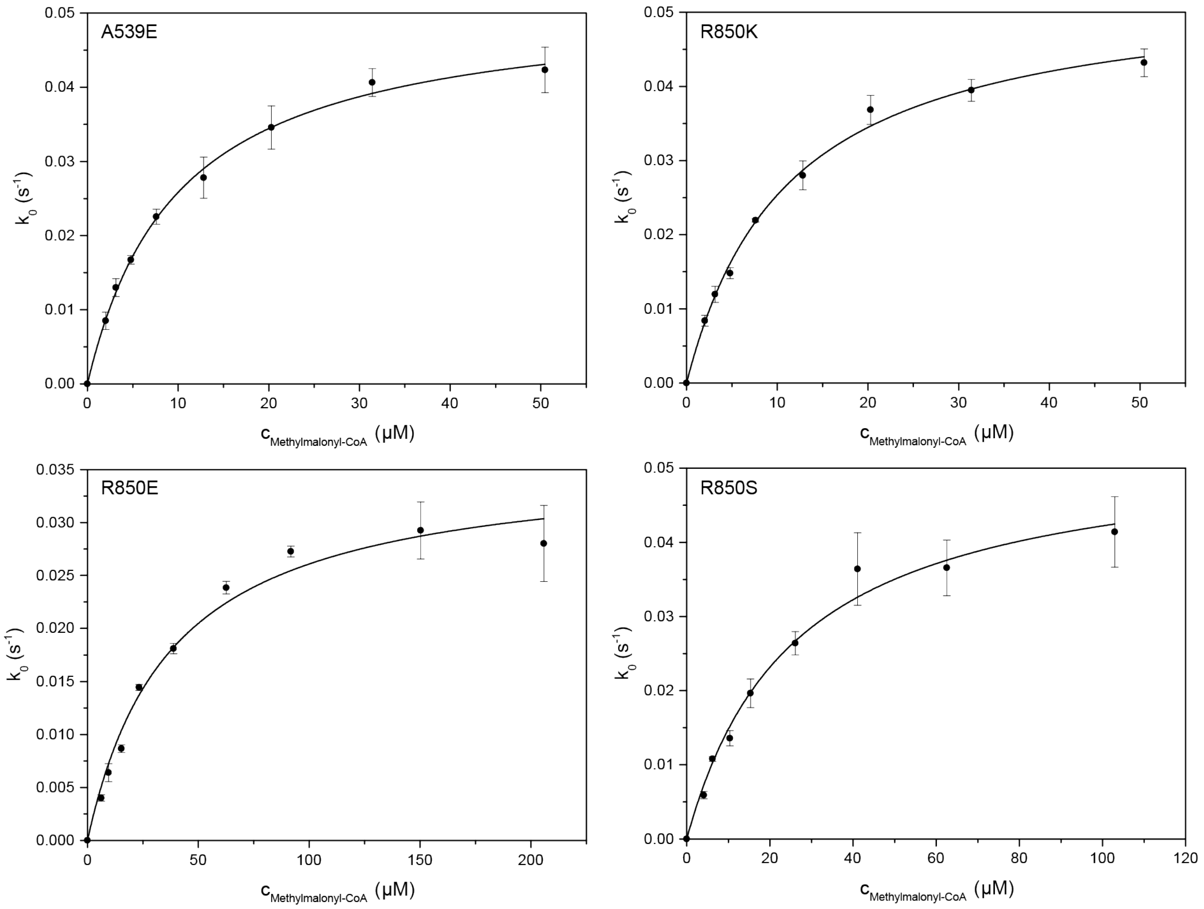

**Figure S12: Titration curves for AT-mediated hydrolysis of MMal-CoA using DEBS3M5 KS$^0$-AT mutants A539E, R850K, R850E and R850S.** Hydrolysis was measured in biological triplicates. Black dots show the average. Error bars correspond to standard deviation of biological triplicates. Kinetic parameters were determined precisely. Substrate consumption for A539E and R850K was below 10 %, for R850E and R850S below 5 %. MMal range was 0.16-5.5 × $K_m^{MMal-CoA}$ (A539E: 0.20-5.1; R850K: 0.18-4.5; R850E: 0.16-5.5; R850S: 0.16-4.0).

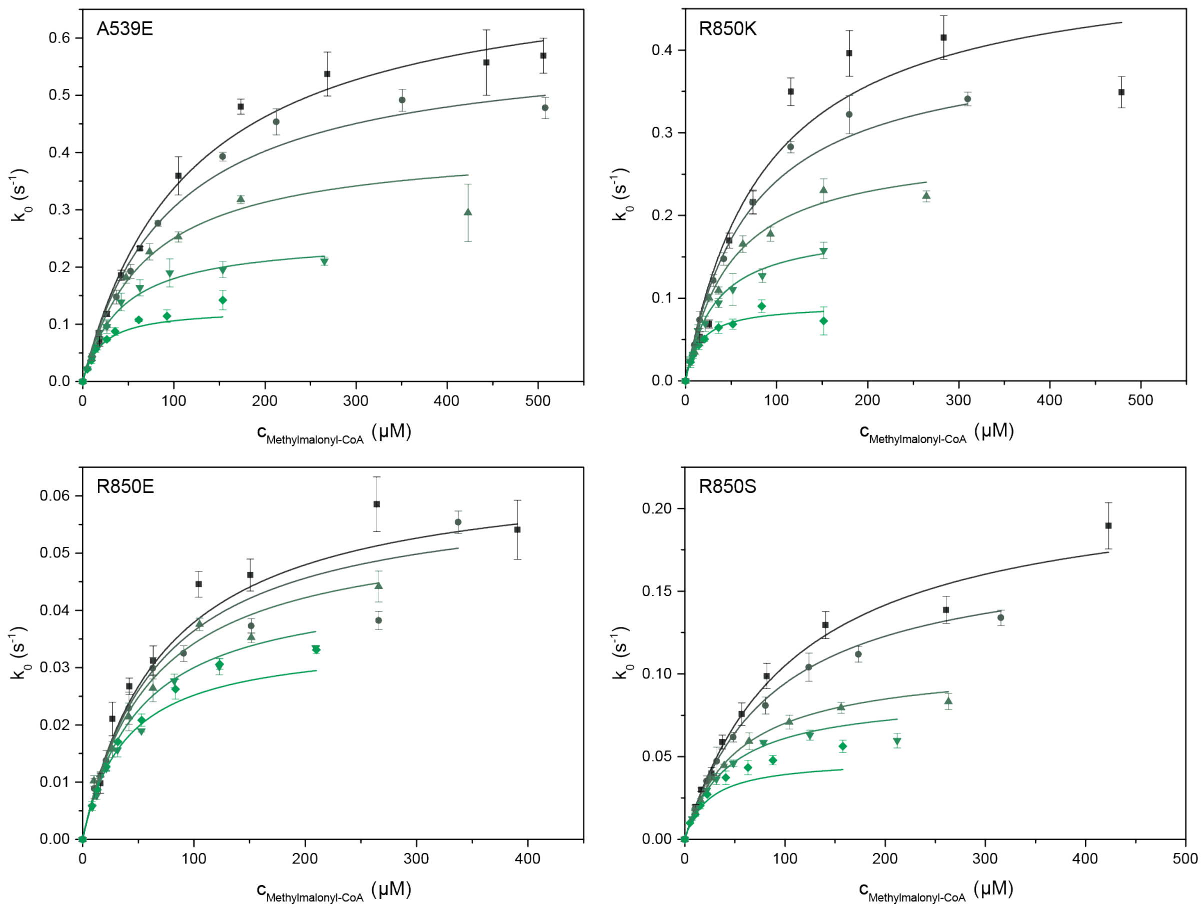

**Figure S13: Global Michaelis-Menten fits of transacylation titration curves with MMal-CoA mediated by DEBS3M5 KS$^{0}$-AT mutants.** All measured in biological triplicates. Each color and symbol corresponds to one X-CoA titration curve at a fixed ACP concentration. Error bars give the standard deviation of biological triplicates. ACP concentrations as follows: A539E: 10.6 µM, 24.3 µM, 51.2 µM, 102.3 µM and 187.3 µM, R850K: 11.0 µM, 26.0 µM, 48.9 µM, 100.4 µM and 190.1 µM, R850E: 2.5 µM, 4.6 µM, 10.0 µM, 20.3 µM and 53.3 µM, R850S: 9.5 µM, 20.5 µM, 28.0 µM, 75.5 µM and 152.4 µM. ACP ranges as follows: A539E: 0.14-2.5 × $K_m^{ACP}$, R850K: 0.15-2.6 × $K_m^{ACP}$, R850E: 1.1-22 × $K_m^{ACP}$, R850S: 0.21-3.3 × $K_m^{ACP}$. Substrate consumption for R850K was below 5 %, for all other proteins was below 10 %. MMal-CoA concentration range as follows: A539E: 0.092-4.9 × $K_{m,app}^{MMal-CoA}$, R850K: 0.13-9.9 × $K_{m,app}^{MMal-CoA}$, R850E: 0.15-6.0 × $K_{m,app}^{MMal-CoA}$, R850S: 0.096-6.6 × $K_{m,app}^{MMal-CoA}$.

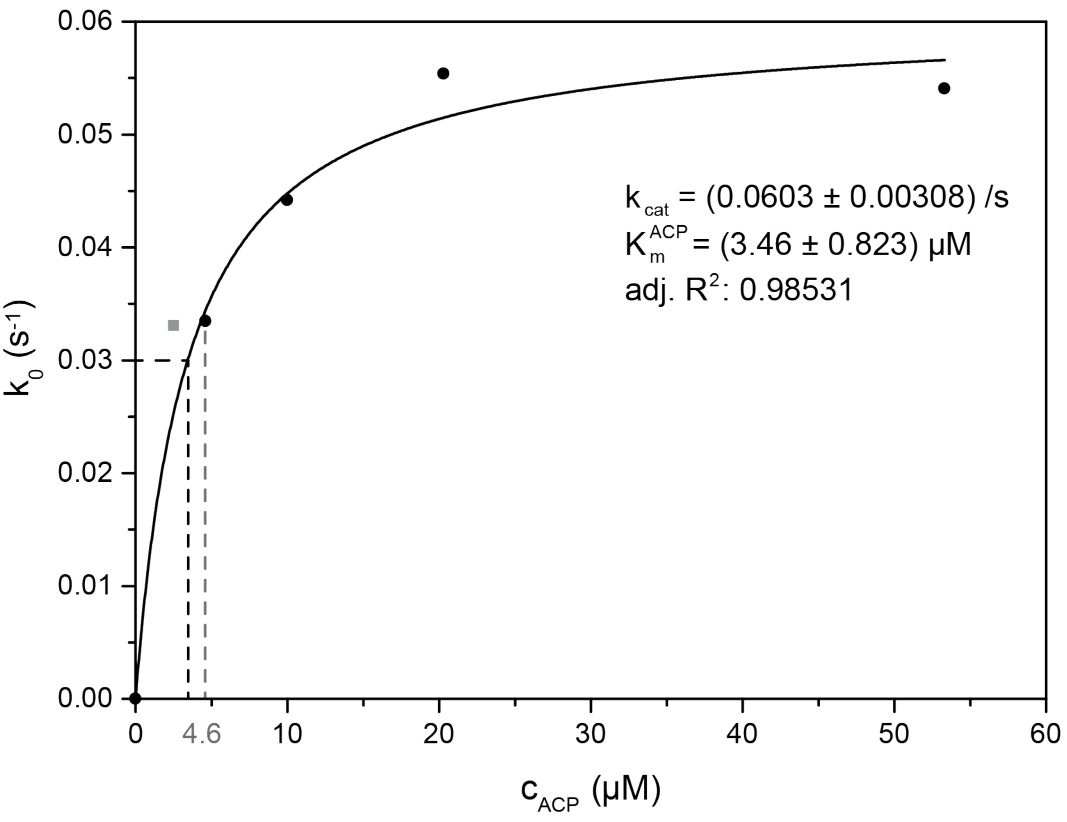

**Figure S14: ACP titration curve at saturated MMal-CoA concentration of mutant R850E.** Due to high measurement errors at low ACP concentrations, the kinetic parameters determined via the global fit are rather error prone. The ACP titration curve is used to determine more reliable kinetic parameters for the AT–ACP interaction. This data was not separately determined, but data points are extracted from MMal-CoA titrations with increasing ACP concentrations (see Figure S13). The lowest ACP concentration (grey box) is omitted to give a more accurate Michaelis-Menten fit. The new plot gives a comparable turnover rate, but a Michaelis-Menten constant of 3.46 µM, which is more reliable than determined before. This value is only 4.8 % of the wild type $K_m^{ACP}$, which is almost the same percentage as determined for the turnover rate $k_{cat}$. The second lowest ACP concentration (4.6 µM) has a $k_0$ above half $k_{cat}$. We have taken this value as upper boundary of the Michaelis-Menten constant.

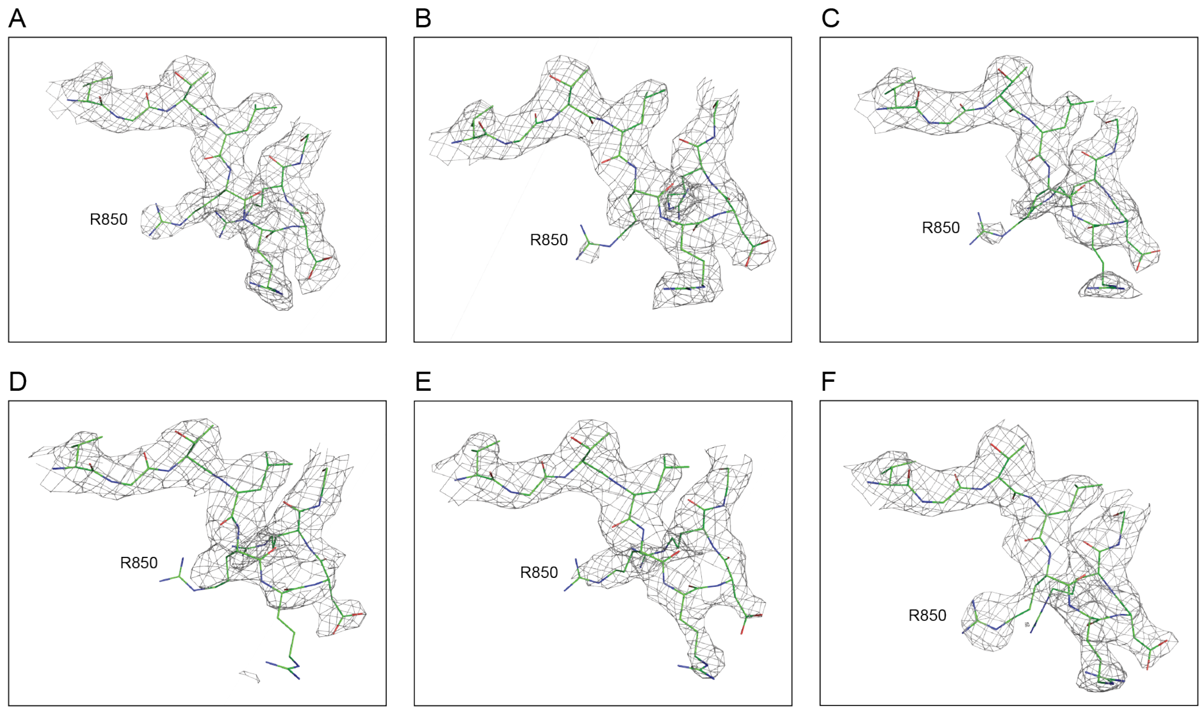

**Figure S15: Electron density map of residue R850.** The electron density map for the residues V846-G854 of DEBS3M5 AT (PDB: 2hg4) for all six chains (A-F) shows poor electron density for the interfacial residue R850 mutated in this study. This finding supports the flexibility of this residue. Different binding modes might be involved during catalysis. The 2fo-fc map was created using PyMOL (MacPyMOL Version 1.7.0.3, Schrödinger, LLC) (sigma=1, carve=1.8).

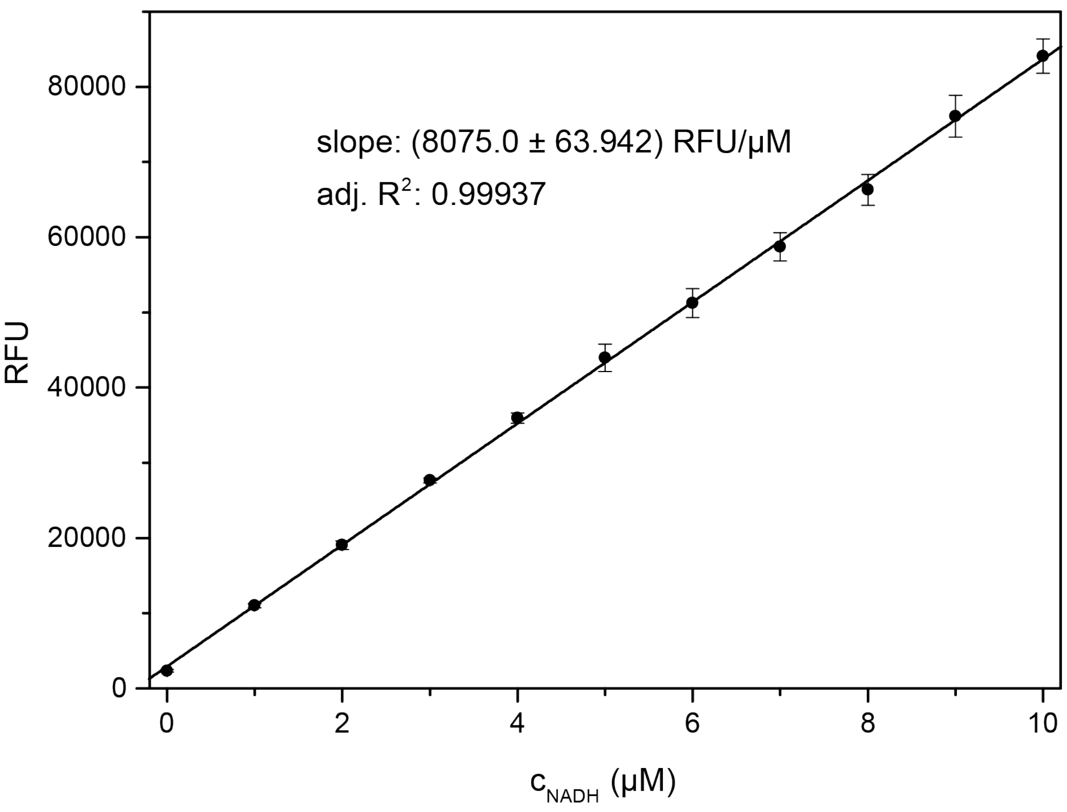

**Figure S16: NADH calibration.** For quantification of AT activity, an NADH calibration was performed. 8075 relative fluorescence units (RFU) correspond to 1 µM NADH. The NADH calibration factor changes over time due to the decreasing power of the UV lamp. An internal standard (50 µM X-CoA and 50 µM ACP) was measured the same day as the NADH calibration. This internal standard was used to determine the calibration factor for each measurement time span. This factor was subsequently used to convert RFU into concentrations (µM) and ranges from 13450 to 8075 RFU per 1 µM NADH. Error bars represent the standard deviation of three independent measurements that were measured in technical triplicates.

### Supporting Information Note 1

#### Derivation of Kinetic Parameters Describing AT-mediated Reactions

Derivation of transacylation parameters as follows. Bound substrate X-CoA, AT-X and X-ACP is abbreviated with XCoA, ATX and XACP.

Notation:

$c_1 \coloneqq [AT] \quad \tilde{c}_1 \coloneqq [XCoA] \quad c_2 \coloneqq [AT \cdot XCoA] \quad c_3 \coloneqq [ATX] \quad \tilde{c}_3 \coloneqq [ACP] \quad c_4 \coloneqq [ATX \cdot ACP]$

$c_{1,0} \coloneqq [AT]_0 \quad \dot{x} \coloneqq \frac{dx}{dt}$

Kinetic equations following steady state assumption:

$I \qquad v \coloneqq \frac{d[XACP]}{dt} = k_4 c_4$

$II \qquad 0 = \dot{c}_4 = k_3 c_3 \tilde{c}_3 - k_{-3} c_4 - k_4 c_4 \qquad \Rightarrow \quad k_3 c_3 \tilde{c}_3 = (k_{-3} + k_4) c_4$

$III \qquad 0 = \dot{c}_3 = k_2 c_2 - k_3 c_3 \tilde{c}_3 + k_{-3} c_4 \qquad \Rightarrow \quad k_3 c_3 \tilde{c}_3 = k_2 c_2 + k_{-3} c_4$

$IV \qquad 0 = \dot{c}_2 = k_1 c_1 \tilde{c}_1 - k_{-1} c_2 - k_2 c_2 \qquad \Rightarrow \quad k_1 c_1 \tilde{c}_1 = (k_{-1} + k_2) c_2$

$V \qquad c_1 = c_{1,0} - c_2 - c_3 - c_4$

Mathematical transformations:

$I = II \qquad \Rightarrow \quad (k_{-3} + k_4) c_4 = k_2 c_2 + k_{-3} c_4 \qquad \Rightarrow \quad c_2 = \frac{k_4}{k_2} c_4$

$c_2$ into $IV$: $IVa \qquad \Rightarrow \quad k_1 c_1 \tilde{c}_1 = (k_{-1} + k_2) \frac{k_4}{k_2} c_4$

$c_2$ into $V$: $Va \qquad \Rightarrow \quad c_1 = c_{1,0} - c_3 - \frac{k_2 + k_4}{k_2} c_4$

$Va$ into $IVa$: $IVb \qquad \Rightarrow \quad k_1 \tilde{c}_1 \left( c_{1,0} - c_3 - \frac{k_2 + k_4}{k_2} c_4 \right) = (k_{-1} + k_2) \frac{k_4}{k_2} c_4$

$\qquad \Rightarrow \quad \tilde{c}_1 c_3 = c_{1,0} \tilde{c}_1 - \left( \frac{k_2 + k_4}{k_2} \tilde{c}_1 + \frac{k_4 (k_{-1} + k_2)}{k_1 k_2} \right) c_4$

$II \times \tilde{c}_1$: $IIa \qquad \Rightarrow \quad k_3 \tilde{c}_1 c_3 \tilde{c}_3 = (k_{-3} + k_4) c_4 \tilde{c}_1$

$IVb$ into $IIa$: $IIb \qquad \Rightarrow \quad k_3 \tilde{c}_3 \left[ c_{1,0} \tilde{c}_1 - \left( \frac{k_2 + k_4}{k_2} \tilde{c}_1 + \frac{k_4 (k_{-1} + k_2)}{k_1 k_2} \right) c_4 \right] = (k_{-3} + k_4) c_4 \tilde{c}_1$

$$\Rightarrow \quad k_3 c_{1,0} \tilde{c}_3 \tilde{c}_1 = \left( \frac{k_3 k_4 (k_{-1} + k_2)}{k_1 k_2} \tilde{c}_3 + \frac{k_3 (k_2 + k_4)}{k_2} \tilde{c}_3 \tilde{c}_1 + (k_{-3} + k_4) \tilde{c}_1 \right) c_4$$

$$= \frac{k_3 (k_2 + k_4)}{k_2} \left( \frac{k_4 (k_{-1} + k_2)}{k_1 (k_2 + k_4)} \tilde{c}_3 + \tilde{c}_3 \tilde{c}_1 + \frac{k_2 (k_{-3} + k_4)}{k_3 (k_2 + k_4)} \tilde{c}_1 \right) c_4$$

$$\Rightarrow \quad c_4 = \frac{\frac{k_2}{k_2 + k_4} c_{1,0} \tilde{c}_3 \tilde{c}_1}{\frac{k_4 (k_{-1} + k_2)}{k_1 (k_2 + k_4)} \tilde{c}_3 + \tilde{c}_3 \tilde{c}_1 + \frac{k_2 (k_{-3} + k_4)}{k_3 (k_2 + k_4)} \tilde{c}_1}$$

$c_4$ into $I$: $$v = \frac{\frac{k_2 k_4}{(k_2+k_4)} c_{1,0} \tilde{c}_3 \tilde{c}_1}{\frac{k_4(k_{-1}+k_2)}{k_1(k_2+k_4)} \tilde{c}_3 + \tilde{c}_3 \tilde{c}_1 + \frac{k_2(k_{-3}+k_4)}{k_3(k_2+k_4)} \tilde{c}_1}$$

For a double displacement reaction the velocity equation is defined as follows:

$v = \frac{v_{max} \tilde{c}_3 \tilde{c}_1}{K_m^{\tilde{c}_1} \tilde{c}_3 + \tilde{c}_3 \tilde{c}_1 + K_m^{\tilde{c}_3} \tilde{c}_1}$ with $v_{max} = k_{cat} c_{1,0}$. [4]

This gives the following kinetic transacylation parameters:

$$k_{cat} = \frac{k_2 k_4}{k_2 + k_4}$$

$$K_m^{XCoA} = \frac{k_4\,(k_{-1} + k_2)}{k_1\,(k_2 + k_4)} = \frac{k_{cat}}{k_1} + \frac{k_{-1} k_4}{k_1\,(k_2 + k_4)}$$

$$K_m^{ACP} = \frac{k_2\,(k_{-3} + k_4)}{k_3\,(k_2 + k_4)} = \frac{k_{cat}}{k_3} + \frac{k_2 k_{-3}}{k_3\,(k_2 + k_4)}$$

$$k_{cat}/K_m^{XCoA} = \frac{k_1 k_2}{k_{-1} + k_2}$$

$$k_{cat}/K_m^{ACP} = \frac{k_3 k_4}{k_{-3} + k_4}$$

Important kinetic parameters describing transacylation are the turnover number $k_{cat}$ and Michaelis-Menten constants $K_m^{X-CoA}$ and $K_m^{ACP}$ for both substrates X-CoA and ACP, respectively. The catalytic efficiency is defined by the ratio of the kinetic parameters $k_{cat}/K_m$. During hydrolysis, only one substrate (X-CoA) is present, simplifying this reaction and leading to less kinetic parameters describing this reaction.

Derivation of hydrolysis parameters as follows. Bound substrate X-CoA, AT-X and X-OH is abbreviated with XCoA, ATX and XOH.

Notation:

$c_1 \coloneqq [AT] \quad \tilde{c}_1 \coloneqq [XCoA] \quad c_2 \coloneqq [AT \cdot XCoA] \quad c_3 \coloneqq [ATX] \quad \tilde{c}_3 \coloneqq [H_2O] \quad c_5 \coloneqq [ATX \cdot OH]$

$c_{1,0} \coloneqq [AT]_0 \quad \dot{x} \coloneqq \frac{dx}{dt}$

The derivation is done analogously, the following parameters change from transacylation to hydrolysis:

$c_4 \rightarrow c_5 \qquad k_3 \rightarrow k_4 \qquad k_4 \rightarrow k_6 \qquad k_{-3} \rightarrow 0$

This gives the following reaction velocity:

$$v = \frac{\frac{k_2 k_6}{(k_2+k_6)} c_{1,0} \tilde{c}_3 \tilde{c}_1}{\frac{k_6(k_{-1}+k_2)}{k_1(k_2+k_6)} \tilde{c}_3 + \tilde{c}_3 \tilde{c}_1 + \frac{k_2 k_6}{k_3(k_2+k_6)} \tilde{c}_1}$$

Setting $\tilde{c}_3 = 1$, since working in aqueous solution, gives v as follows:

$$v = \frac{\frac{k_2 k_6}{(k_2+k_6)} c_{1,0} \tilde{c}_1}{\frac{k_6(k_{-1}+k_2)}{k_1(k_2+k_6)} + \left(1 + \frac{k_2 k_6}{k_3(k_2+k_6)}\right) \tilde{c}_1} = \frac{\frac{k_2 k_5 k_6}{k_2 k_6 + k_5(k_2+k_6)} c_{1,0} \tilde{c}_1}{\frac{k_5 k_6 (k_{-1}+k_2)}{k_1\left(k_2 k_6 + k_5(k_2+k_6)\right)} + \tilde{c}_1}$$

This gives the following kinetic hydrolysis parameters. Notably the catalytic efficiency $k_{cat}/K_m^{XCoA}$ is the same for transacylation and hydrolysis.

$$k_{cat} = \frac{k_2 k_5 k_6}{k_2 k_6 + k_5(k_2 + k_6)}$$

$$K_m^{XCoA} = \frac{k_5 k_6 (k_{-1} + k_2)}{k_1\left(k_2 k_6 + k_5(k_2 + k_6)\right)} = \frac{k_{cat}}{k_1} + \frac{k_{-1} k_5 k_6}{k_1\left(k_2 k_6 + k_5(k_2 + k_6)\right)}$$

$$k_{cat}/K_m^{XCoA} = \frac{k_1 k_2}{k_{-1} + k_2}$$

### Supporting Information Material and Methods

#### Material and Methods

List of expression plasmids used in this study with protein mass and absorbance.

Protein size given for proteins without N-terminal methionine and for phosphopantetheinylated ACP. Protein absorbance values (Abs) calculated with CLC Main Workbench.

| # | Plasmid Name | Protein Size (kDa) | Abs (g/L) |
|---|---|---|---|
| pAR001 | SFP_bsub_pETcoco | | |
| pAR225 | T7_StrepI_Pks7(KS(C161G)_AT)_H8_pET22b | 98.552 | 1.266 |
| pAR331 | T7_StrepI_DEBS3(ACP)5_H8_pET22b | 15.375 | 0.842 |
| pAR333 | T7_StrepI_RAPS3(ACP)14_H8_pET22b | 14.144 | 0.917 |
| pFS001 | T7_StrepI_DEBS3M5(KS(C233G)_AT)_H8_pET22b | 99.850 | 1.021 |
| pFS004 | T7_StrepI_PikAIIIM5(KS(C243G)_AT)_H8_pET22b | 100.679 | 1.124 |
| pFS007 | T7_StrepI_RAPS3M14(KS(C226G)_AT)_H8_pET22b | 100.504 | 1.208 |
| pFS120 | T7_StrepI_DEBS3(ACP)5-COP_H8_pET22b | 15.375 | 0.842 |
| pFS121 | T7_StrepI_PikAIII(ACP)5-COP_H8_pET22b | 15.177 | 0.857 |
| pFS123 | T7_StrepI_Pks7(ACP)-COP_H8_pET22b | 16.366 | 1.500 |
| pFS129 | T7_StrepI_DEBS3M5(KS(C233G)_AT(A539S))_H8_pET22b | 99.866 | 1.021 |
| pFS130 | T7_StrepI_DEBS3M5(KS(C233G)_AT(A539D))_H8_pET22b | 99.894 | 1.021 |
| pFS131 | T7_StrepI_DEBS3M5(KS(C233G)_AT(A539E))_H8_pET22b | 99.908 | 1.021 |
| pFS132 | T7_StrepI_DEBS3M5(KS(C233G)_AT(A539F))_H8_pET22b | 99.926 | 1.021 |
| pFS133 | T7_StrepI_DEBS3M5(KS(C233G)_AT(R850K))_H8_pET22b | 99.822 | 1.022 |
| pFS134 | T7_StrepI_DEBS3M5(KS(C233G)_AT(R850A))_H8_pET22b | 99.765 | 1.022 |
| pFS135 | T7_StrepI_DEBS3M5(KS(C233G)_AT(R850E))_H8_pET22b | 99.823 | 1.022 |
| pFS136 | T7_StrepI_DEBS3M5(KS(C233G)_AT(R850F))_H8_pET22b | 99.841 | 1.022 |
| pFS137 | T7_StrepI_DEBS3M5(KS(C233G)_AT(R850S))_H8_pET22b | 99.781 | 1.022 |

#### Protein Expression

All constructs were transformed into *E. coli* BL21gold (DE3) cells (Agilent Technologies) following the protocol of the manufacturer. All vectors encoding ACPs were co-transformed with Sfp. LB agar (Lennox) transformation plates were supplemented with 1 % glucose and the respective antibiotics. As pre-culture, 20 mL Lysogeny Broth medium supplemented with 1 % glucose and the respective antibiotics were inoculated with one single clone and incubated at 37 °C and 180 rpm overnight. 1 L Terrific Broth medium supplemented with the respective antibiotics was inoculated with the pre-culture and incubated at 37 °C and 130-

180 rpm until an $OD_{600}$ of 0.6-0.8 was reached. After cooling for 15 min at 4 °C, expression of proteins was induced with 0.25 mM IPTG final concentration. All proteins were expressed at 20 °C and 130-180 rpm overnight for approximately 16 h. Cells were harvested by centrifugation at 4,000 rpm for 20 min. Cell pellets were immediately purified or stored at -80 °C for several days.

Protein Purification

Cell pellets were resuspended in corresponding Ni Buffer (AT: 450 mM NaCl, 50 mM $NaH_2PO_4$, 10 mM imidazole, 20 % v/v glycerol, pH 7.6; ACP: 200 mM NaCl, 50 mM $NaH_2PO_4$, 20 mM imidazole, 10 % v/v glycerol, pH 7.4) containing DNAse I and 1 mM EDTA. Cells were mechanically disrupted at 1,000 bar using a French pressure cell press and centrifuged at 50,000 rcf and 4 °C for 1 h. After centrifugation, 2 mM $MgCl_2$ were added. All AT constructs were purified by tandem affinity chromatography (Ni-NTA and Strep-Tactin), all ACPs were purified by Ni-affinity chromatography. Lysate was loaded onto 5 mL Ni-NTA columns, unbound protein was washed off with 5 CV Ni Buffer. Bound protein was eluted with 2.5 CV Ni Buffer containing 300 mM imidazole. For AT constructs, Ni elution fractions were loaded onto 5 mL Strep-Tactin columns (Iba), unbound protein was washed off with 6 CV Strep Buffer (500 mM NaCl, 50 mM $NaH_2PO_4$, 20 % v/v glycerol, pH 7.6). Bound protein was eluted with 2.5 CV Strep Buffer containing 2.5 mM desthiobiotin. All ACPs were further purified by size exclusion chromatography (SEC) using HiLoad 16/600 Superdex 200 pg column (200 mM NaCl, 50 mM $NaH_2PO_4$, 10 % v/v glycerol, pH 7.4, filtered and degased). Elution fractions were analyzed by SDS-PAGE. ACP was pooled after SEC to create one stock of each ACP as substrate used for all measurements of the AT activity assay. This should minimize errors in assay resulting from inconsistent protein expression/quality. Proteins were concentrated using Amicon Ultra Centrifugal Filters (Merck Millipore), flash frozen in liquid nitrogen and stored in aliquots at -80 °C.

Protein Quality Control of AT Constructs

Quality and oligomeric state of all AT constructs were analyzed by HPLC-SEC (500 mM NaCl, 50 mM $NaH_2PO_4$, 5 % v/v glycerol, pH 7.6) at room temperature with a flow of 0.3 ml/min. An UltiMate 3000 RSLC (Dionex) equipped with an UV-vis array detector was used with SEC columns Yarra-SEC-X150 and Yarra-SEC-X300. Proteins were detected by absorbance at 280 nm. A thermal shift assay (TSA) was used to analyze all AT constructs using CFX96 Touch Real-Time PCR Detection System (BioRad) with excitation and emission wavelength set to 450-490 and 560-580 nm, respectively. Fluorescence of the dye SYPRO Orange was measured from 5 to 95 °C with 0.5 °C steps per minute in Multiplate 96-well

PCR Plates (BioRad). Melting curves of all wild type AT constructs were measured in storage buffer and AT activity assay buffer. DEBS3M5 AT mutants were measured in storage buffer. Analysis: OriginPro 9.1.

##### Mass Spectrometric Analysis of ACPs

Full phosphopantetheinylation of ACPs was controlled by mass spectrometry. Proteins were precipitated and resolved in sterile water. An ultimate 3000 RSLC system (Dionex) coupled to an impact II (Bruker, for non-codon optimized ACPs) or to a micrOTOF-Q II (Bruker, for codon optimized ACPs) equipped with an electrospray ionization source was used to perform HPLC-MS of ACPs. For chromatographic separation an RP-C3 column (Zorbax 300-SB, 300 Å, 150 mm × 3.0 mm × 3.5 µm, Agilent) was used with a mobile phase system consisting of solvent A (water with 0.1 % (v/v) formic acid) and solvent B (acetonitrile with 0.1 % (v/v) formic acid). After equilibration with 15 % solvent B and a flow of 0.6 mL/min for 1.5 min, the concentration of solvent B was linearly increased to 65 % over 22.5 min. This was followed by a linear increase of solvent B to 95 % over 4 min prior to re-equilibration of 15 % solvent B for 2 min. Proteins were detected by absorbance at 280 nm. ACPs were found to elute at 9.4-16 min. MS data was acquired in positive mode in the range of 50-2000 m/z and analyzed using DataAnalysis 4.3 software (Bruker Dalton GmbH).

##### AT Activity Assay

The AT activity assay was adapted from references.[1], [5], [6] The AT activity assay buffer (50 mM $NaH_2PO_4$, 1 mM EDTA, 1 mM DTT added directly before usage, 10 % v/v glycerol, pH 7.6) was used to set up all other solutions for this assay. For all proteins and substrates, fresh aliquots were used. For each system, the same batch of substrates (ACP and acyl-CoAs) was used. Four master mixes in 4 × concentration were prepared. For the acyltransferase solutions the proteins were diluted to 0.1-400 nM final assay concentration with assay buffer supplemented with 0.1 mg/mL BSA. The ACP solution was prepared by diluting highly pure and concentrated ACP to 2.5-530 µM final concentration with the assay buffer. Substrate acyl-CoA solution was also prepared with the assay buffer in different concentrations ranging from 1 µM to 1 mM final concentration. The read out solution contained 2 mM α-ketoglutaric acid, 0.4 mM $NAD^+$, 0.4 mM TPP and 5 to 15 mU/100 µL αKGDH as final concentrations. All solutions were pre-heated/incubated at 25 °C for at least 5 min. Assays were performed in 96-well f-bottom microtiter plates (Greiner Bio-one). For transacylation, 25 µL of each solution was added, ACP was added via the dispenser to initiate the reaction. For hydrolysis, assay buffer was used to compensate the volume of

ACP. For background measurements the AT solution was replaced by 0.1 mg/mL BSA in assay buffer. NADH fluorescence was monitored for 5 min at 25 °C, taking equidistant measurements every 20 s using a ClarioStar microplate reader (BMG labtech). The following settings were used: excitation: 340-20 nm, emission: 476-20 nm, gain: 1900, focal height: 5.2 mm, flashes: 70, orbital averaging: 4 mm.

To meet Michaelis-Menten requirements, the substrate concentration was always kept significantly higher than the enzyme concentration, at least ten times. [4] For hydrolysis and for each ACP curve in transacylation measurements, initial experiments were performed to estimate the (apparent) $K_m^{X-CoA}$. Based on these, the acyl-CoA substrate concentration was varied to calculate kinetic parameters: (0.2; 0.3; 0.5; 0.75; 1.25; 2; 3; 5) × $K_{m(,app)}^{X-CoA}$. In order to ensure that the measurement is performed in the initial quasi-linear range of substrate consumption, the product concentration has to be significantly lower than the substrate concentration. [4] The increase during measurement was always linear and substrate consumption was ensured to be below 10 % for the majority of assays performed. Some measurements showed significantly higher substrate consumption. Nevertheless, it was always ensured to be in a linear range and the global Michaelis-Menten fit is robust and the kinetic parameters do not change, if these values are omitted. If not mentioned otherwise, transacylation was performed in biological triplicates. Hydrolysis was performed in technical triplicates of biological triplicates or technical/biological triplicates. Background values were subtracted from the average of technical/biological measurement values. Occasional dust or air bubbles caused strong outliers. These were omitted for further analysis after careful analysis. The time span of 40-120 and 20-120 s was analyzed for transacylation and hydrolysis, respectively. For high substrate consumption, this time span was adapted. In order to convert RFU into concentration, an NADH calibration curve was measured (**Figure S14**). Hydrolysis kinetic parameters were calculated by Michaelis-Menten plots using OriginPro 9.1. Transacylation parameters were calculated by global Michaelis-Menten fits using OriginPro 9.1.

##### Rational Design of Mutations in the AT:ACP Interface

Since the AT:ACP interface of DEBS3M5 and the structure of its ACP is not known, SwissModel was used to model the ACP domain. [7] Subsequently, three AT:ACP interfaces were used as template for structural alignment of the modeled ACP and the structure of AT domain (PDB: 2hg4). As templates served the *in silico* docking study on human FAS and crystal structures of the AT:ACP complex of vicenistatin PKS VinK:VinL and of disorazole synthase (DSZS). [8–10] Based on these three alignments, PISA was used to identify hotspots in the AT:ACP interface.[11] All three models gave the amino acids A539 and R850 as possible interfacial hotspots.

#### Amino Acid Sequences

DEBS3M5 ACP

MSAWSHPQFEKGGGSGGGSGGSAWSHPQFEKGAGSGPALAQRLAALSTAERREHLAHLI
RAEVAAVLGHGDDAAIDRDRAFRDLGFDSMTAVDLRNRLAAVTGVREAATVVFDHPTITRLA
DHYLERLVGALEHHHHHHHH

PikAIIIM5 ACP

MSAWSHPQFEKGGGSGGGSGGSAWSHPQFEKGAGSGQSSALAAITALPEPERRPALLTL
VRTHAAAVLGHSSPDRVAPGRAFTELGFDSLTAVQLRNQLSTVVGNRLPATTVFDHPTPAA
LAAHLHEAYLAPLEHHHHHHHH

RAPS3M14 ACP

MSAWSHPQFEKGGGSGGGSGGSAWSHPQFEKGAGSPSYADEPRTMLELVHMEVASLLG
MADPGVILDDSSFLELGFDSLSAVRLRNRLSKATGLDLPSTLLFEHPTSAELAAHLDALLDSL
EHHHHHHHH

Pks7 ACP

MSAWSHPQFEKGGGSGGGSGGSAWSHPQFEKGAGSEADAGPAEYRHLDPERLHEWLTV
EVVKLIAAEMRLHADELDPTASLVTQGLDSVMSLVLRRRLEKRFGQSLPANLLWHKPTAAAI
VDHLSGLLSDNVREGAASLEHHHHHHHH

DEBS3M5 KS$^0$-AT

MSAWSHPQFEKGGGSGGGSGGSAWSHPQFEKGAGSSGDNGMTEEKLRRYLKRTVTELD
SVTARLREVEHRAGEPIAIVGMACRFPGDVDSPESFWEFVSGGGDAIAEAPADRGWEPDP
DARLGGMLAAAGDFDAGFFGISPREALAMDPQQRIMLEISWEALERAGHDPVSLRGSATGV
FTGVGTVDYGPRPDEAPDEVLGYVGTGTASSVASGRVAYCLGLEGPAMTVDTAGSSGLTA
LHLAMESLRRDECGLALAGGVTVMSSPGAFTEFRSQGGLAADGRCKPFSKAADGFGLAEG
AGVLVLQRLSAARREGRPVLAVLRGSAVNQDGASNGLTAPSGPAQQRVIRRALENAGVRA
GDVDYVEAHGTGTRLGDPIEVHALLSTYGAERDPDDPLWIGSVKSNIGHTQAAAGVAGVMK
AVLALRHGEMPRTLHFDEPSPQIEWDLGAVSVVSQARSWPAGERPRRAGVSSFGISGTNA
HVIVEEAPEADEPEPAPDSGPVPLVLSGRDEQAMRAQAGRLADHLAREPRNSLRDTGFTLA
TRRSAWEHRAVVVGDRDDALAGLRAVADGRI**A**DRTATGQARTRRGVAMVFPGQGAQWQ
GMARDLLRESQVFADSIRDCERALAPHVDWSLTDLLSGARPLDRVDVVQPALFAVMVSLAA
LWRSHGVEPAAVVGHSQGEIAAAHVAGALTLEDAAKLVAVRSRVLRRLGGQGGMASFGLG
TEQAAERIGRFAGALSIASVNGPRSVVVAGESGPLDELIAECEAEGITARRIPVDYASHSPQV
ESLREELLTELAGISPVSADVALYSTTTGQPIDTATMDTAYWYANLREQVRFQDATRQLAEA
GFDAFVEVSPHPVLTVGIEATLDSALPADAGACVVGTL**R**RDRGGLADFHTALGEAYAQGVE
VDWSPAFADARPVELPVYPFQRQRYWLPIPTGGLEHHHHHHHH

PikAIIIM5 KS$^0$-AT

MSAWSHPQFEKGGGSGGGSGGSAWSHPQFEKGAGSANNEDKLRDYLKRVTAELQQNTRRLREIEGRTHEPVAIVGMACRLPGGVASPEDLWQLVAGDGDAISEFPQDRGWDVEGLYDPDPDASGRTYCRSGGFLHDAGEFDADFFGISPREALAMDPQQRLSLTTAWEAIESAGIDPTALKGSGLGVFVGGWHTGYTSGQTTAVQSPELEGHLVSGAALGFLSGRIAYVLGTDGPALTVDTAGSSSLVALHLAVQALRKGECDMALAGGVTVMPNADLFVQFSRQRGLAADGRSKAFATSADGFGPAEGAGVLLVERLSDARRNGHRILAVVRGSAVNQDGASNGLTAPHGPSQQRVIRRALADARLAPGDVDVVEAHGTGTRLGDPIEAQALIATYGQEKSSEQPLRLGALKSNIGHTQAAAGVAGVIKMVQAMRHGLLPKTLHVDEPSDQIDWSAGTVELLTEAVDWPEKQDGGLRRAAVSSFGISGTNAHVVLEEAPAVEDSPAVEPPAGGGVVPWPVSAKTPAALDAQIGQLAAYADGRTDVDPAVAARALVDSRTAMEHRAVAVGDSREALRDALRMPEGLVRGTSSDVGRVAFVFPGQGTQWAGMGAELLDSSPEFAASMAECETALSRYVDWSLEAVVRQEPGAPTLDRVDVVQPVTFAVMVSLAKVWQHHGITPQAVVGHSQGEIAAAYVAGALTLDDAARVVTLRSKSIAAHLAGKGGMISLALDEAAVLKRLSDFDGLSVAAVNGPTATVVSGDPTQIEELARTCEADGVRARIIPVDYASHSRQVEIIEKELAEVLAGLAPQAPHVPFFSTLEGTWITEPVLDGTYWYRNLRHRVGFAPAVETLAVDGFTHFIEVSAHPVLTMTLPETVTGLGTLRREQGGQERLVTSLAEAWANGLTIDWAPILPTATGHHPELPTYAFQTERFWLQSSAPTSAALEHHHHHHHH

RAPS3M14 KS$^0$-AT

MSAWSHPQFEKGGGSGGGSGGSAWSHPQFEKGAGSEAPAPSTVTGPADPVADEPSANEPIAIVAMACRLPGGVSSPEGLWHLVESGTDAISGFPTDRGWDVEGLFDPDPDAAGKSYCVQGGFLDTAADFDAPFFGISPREALGMDPQQRLLLETTWEAIERAQIDPKSLRGRDVGVYVGGAAQGYGVGVDQQHDNGITGSSVSLLSGRVSYALGLEGPGVTVDTAGSSSLVALHLASQALRQRECSLALVSGVSVMSSPAMFVEFSRQRGLSSDGRCKSFAASADGTIWSEGVGVLVVERLSDARRLGHRVLATVRGSAVNSDGASNGLTAPNGTSQQRVIRQALANAGLTASDVDVVEAHGTGTKLGDPIEAEAILATYGQERSAPAWLGSLKSNIGHAMAASGVLSVIKMVEAMAHGSLPRTLHVDAPSPHVDWTSGSVALLTEHQPWPDDTKLRRAGVSSFGLSGTNAHVVLEQYQAPAPPVTPVTPAPPVTPVTPVTPNEPGPLPWVLSAQSPKALREQAGRLYASLAGDSEWNSLDIGYSLATTRSDFAHRAVAVGSGREDFLRALSKLADGAPWPGLTTATATAKARRVAFLFDGQGTQRLGMGKELYDSYPAFARAWDTVSAGFDKHLDHSLTDVCFGEGGSTTAGLVDDTLYAQAGIFAMEAALFGLLEDWGVRPDFVAGHSIGEATAAYASGMLSLENVTTLIVARGRALRTTPPGAMVALRAGEEEVREFLSRTGAALDLAAVNSPEAVVVSGEPEPVADFEAAWTASGREARKLKVRHAFHSRHVEAVLDEFRTALESLKFRAPALPVVSTVTGRLIDQDEMGTPEYWLRQVRRPVRFQDAVRELAEQGVGTFVEVGPSGALASAGVECLGGDASFHAVLRPRSPEDVCLMTAIAELHAGGTAIDWAKVLSGGRAVDLPVYPFQHQSYWLAPAAPDATALEHHHHHHHH

Pks7 KS$^0$-AT

MSAWSHPQFEKGGGSGGGSGGSAWSHPQFEKGAGSSTQPDAPEPIAVVGIGCRLPGSANSADALWDLLAAGRDVVGEVPDDRWRDYLAMGPAYAAATRRTPRRGGYLDEDIAWFDNEF

FGVTHREAETMDPQHRMMLEVTWEALEHAGIPPLTLAGRQVGVYTGVINDDYGRRLLENLP
DLDAWVAIGVANSGAANRVSYALDLRGPSLAVDTAGSASLTAVHLACQSLRAGESELALAG
GVQLIAAPAWSLSLEAGGFLSPVGLSKAFSGDADGYVRGEGCGVLVLKRLADAERDGDRIL
SVVLGTSVIHDGRSENFVAPSEAAQQAMARQACAEAGIEPNTVDYVEAHGTGTRRGDRTE
VVALSTVYGAGRPADDPCLIGSVKTNIGHLEASAGIAGVIKAVLALGHDHIPPSLHNSSLNPA
VDWETAGIKVVTEPTAWPRRSHPRRAAVSSFGFGGTISHVLLQQAPDRRPAAAVPSSTTPN
TNTSTVFPLSARSESGLRRNAARLADWLRGPGADTALADVGHTLALRRSHLTHRASVVAAD
RDELINGLRNIADGELAPGIATGTTDAVPRAVWVFSGHGSQWAGMGRELLNAQPVFAGVID
KIEPVFAEESGFSLREALREGEFAGVGRTQMLIFAMQVALAEVWRAHGAAPDAVIGHSMGEI
AAAVVSGACTLEVGARLISRRSGLLWRAEGKGTMATINLPFDEVATKLAGRDDVVAAISSSP
RASVISGDVGAVESLVGDWEAAGLLPRRINIQMASHSPAMDPLLADLREAIADLPVGEPLIP
MYSTSATDPRSKGVLDGEYWVGNLRRPVRLREAVEAAVADGYGAFLEVAPHPVVGNPISE
TVSALGREDVFVGLTLRRGHPEHETLMGSIGEAHCHGIDVDLGCLYLEGELATLPSMQWLR
QPHWRDFAPSGALEHHHHHHHH

### Supporting Information References
